# GPCR-MAPS: high-resolution functional and allosteric mapping of G protein-coupled receptor activation and bias

**DOI:** 10.1101/2025.05.30.656974

**Authors:** Taylor L. Mighell, Ben Lehner

## Abstract

G protein-coupled receptors (GPCRs) are the largest family of human receptors and drug targets, but how the signaling properties of receptors are encoded in their sequence is incompletely understood. Most GPCR drugs bind a conserved orthosteric pocket, which can result in non-specificity, toxicity and clinical failure. Precision modalities including biased signaling and allosteric modulation could lead to GPCR therapeutics with improved efficacy and safety profiles. The rational design of precision therapies is, however, limited by the lack of high-resolution functional annotation of GPCR structures. Here we present a fast and general approach to rapidly build complete, high-resolution, multi-modal functional and allosteric maps of receptors. The approach, GPCR-MAPS, quantifies direct recruitment of proteins to a receptor and deconvolves the substantial effects of mutations on receptor expression. Applying GPCR-MAPS to the β2 adrenergic receptor generates >150,000 phenotypic measurements, including the full activation functions for all possible amino acid substitutions (>7,500 unique variants) in a single experiment. The multi-modal maps provide numerous mechanistic insights and reveal a modular receptor architecture, with a core activation network surrounded by residues controlling quantitative parameters and bias. The maps also identify multiple allosteric surface pockets, including pockets bound by serendipitously discovered allosteric ligands and a novel pocket with no known ligands. The application of this approach across the superfamily of GPCRs will provide comparative maps of receptor mechanisms and a functional framework for the rational design of precision GPCR therapeutics.

## Introduction

G-protein coupled receptors (GPCRs) are the largest family of transmembrane proteins that receive and transmit signals from the outside to the inside of cells. These signals are very diverse and include neurotransmitters, metabolites, immune system molecules, pH, and photons of light^1^. The topology of GPCRs is highly conserved and consists of seven transmembrane (TM) helices connected by alternating intra- and extracellular loops (ECLs and ICLs). Binding of an agonist to the extracellular portion of the receptor results in an allosteric conformational change to the intracellular portion of the receptor^2^, which enables binding and activation of one or more members of the family of heterotrimeric G-proteins, which subsequently activate diverse downstream signaling pathways^3^. Receptor activation also results in phosphorylation of the receptor, including in the C-terminal tail, and recruitment of β-arrestin, which terminates G-protein signaling by internalizing the receptor, and also scaffolds G-protein independent signaling^4^. Because of their role as master signal transducers and their availability on the cell surface, GPCRs are the most important class of drug targets, with one third of current clinical therapeutics targeting GPCRs^5,6^.

The vast majority of established GPCR drugs act by binding to the orthosteric site and activating or inactivating the receptor^7^. However, alternative modalities for modulating GPCRs exist, such as biased agonism^8^ (also called “functional selectivity”) and allosteric modulation^9^. Biased agonists are defined in opposition to balanced agonists; a biased agonist preferentially signals through one downstream pathway over another (e.g. G-protein over β-arrestin), compared to balanced agonists. In cases where signaling through one pathway leads to an unwanted side effect, biased agonists may achieve improved therapeutic efficacy. A therapeutic benefit for biased agonism has been proposed for at least thirty GPCRs^10^.

Allosteric drugs are those that bind the receptor at a spatially distinct site from the orthosteric site. The high structural conservation of orthosteric sites makes it difficult to target one receptor without also engaging multiple family members. This non-specificity can result in toxicity and other side-effects that result in clinical failure^7^. In contrast, allosteric drugs targeting non-conserved sites can have high specificity and reduced toxicity. Further, since allosteric drugs modulate responses to physiological ligands, their effects temporally and spatially match physiological signaling processes^11^. Due to these major advantages, increasing effort is being put toward development of biased and allosteric modulators^12^. However, with our current understanding of GPCR function, these molecules are typically only found by blind screening or iteration upon previously discovered molecules^13,14^.

In order to design better GPCR drugs, an improved understanding of the rich functional and allosteric architecture linking agonist binding with intracellular signaling is required. Much work has focused on using the structures of active and inactive states of receptors to try and understand this allosteric mechanism^15,16^. This has provided important mechanistic insights, but these structures are only the starting point for assigning a functional role for the hundreds of individual amino acids in each highly dynamic receptor.

A powerful complementary approach for understanding how proteins work is the use of mutagenesis followed by functional characterization^17^. Serial experiments have measured the effects of hundreds of alanine substitutions on GPCR signaling through one^18^ or two^19,20^ pathways, revealing a rich diversity of signaling responses to individual mutations. Pooled approaches have been used to characterize surface expression^21–25^, ligand binding^23,26^, and single or few dose transcriptional responses downstream of GPCRs^27–30^. However, the limited throughput^18–20^ and lack of quantification of signaling parameters limit the insights that can be obtained by these approaches^21–30^.

Many of the most basic questions in GPCR signaling remain unanswered. Challenges include describing and understanding how the sequence of a receptor determines signaling properties and the causal mapping from sequence to biophysics to signaling characteristics. Properties to understand include cellular abundance, maximum response, basal signaling, and potency of response of a receptor to ligand binding. Mutating receptor residues to alanine has provided important insights into this, but testing only 1 of 19 alternative amino acids in each position provides a limited view of the functional consequences of sequence change. An additional challenge is to understand how the sequence of a receptor controls bias in the recruitment of different transducer proteins^31^.

With respect to drug discovery, a key challenge is to understand how ligands modulate receptor signaling. Are all ligand-contacting sites equally important for a response or just a subset of “hotspot”^32^ residues? Are all ligand contacts equally important for different functional responses to a ligand or are some contacts more important for particular responses? Where else in the receptor mediates the response to a ligand? Are residues important for the response to ligand equally important for different functional responses, including signaling bias?

A major goal in GPCR drug discovery is to identify allosteric sites on a receptor where perturbations can inhibit, activate, or modulate signaling. GPCRs, like all proteins, have many potential small pockets to therapeutically target^33^. For each receptor it would be extremely useful to know at which of these sites perturbations alter receptor activity and how, including quantifying changes in agonist responses and altered bias.

Here we present a general high-throughput method, GPCR-MAPS (GPCR <u>m</u>ultiplexed <u>a</u>ssays with <u>p</u>athway <u>s</u>pecificity), to rapidly generate comprehensive functional maps of GPCRs. As a first application, we apply GPCR-MAPS to the beta-2 adrenergic receptor (β2AR), one of the most extensively characterized receptors. β2AR is expressed in the cardiovascular system where it modulates smooth muscle relaxation and bronchodilation; β2AR agonists are therefore used for treating asthma as well as chronic obstructive pulmonary disease^34^.

GPCR-MAPS has four components: First, we generate a complete site-saturation library of a receptor, consisting of all possible amino acid substitutions in all positions. Second, we quantify the cell surface expression of the library of variants to quantify defects in fold stability and trafficking. Third, we use pooled experiments to quantify the recruitment of conformation-specific interaction partners to every receptor mutant in response to ligand treatment. Using ligand titrations allows the complete dose-response activation functions of thousands of receptor variants to be quantified. Measuring the recruitment of different interaction partners quantifies changes in receptor bias. Fourth, we integrate the resulting data to generate rich multi-modal functional maps of how the quantitative function of a receptor is altered by diverse perturbations throughout its entire sequence.

Applied to β2AR, GPCR-MAPS quantifies the activation functions and multi-modal characteristics of >7,500 different protein variants in pooled experiments. The resulting multi-modal maps reveal a modular receptor architecture, with a core activation network surrounded by residues controlling quantitative parameters and bias. The multi-modal maps also identify multiple allosteric surface pockets, including pockets bound by serendipitously discovered β2AR allosteric ligands and a novel pocket with no known ligands. We believe application of the GPCR-MAPS approach across the GPCR superfamily will advance our understanding of the largest family of human membrane receptors and enable rational, precision drug design.

## Results

### GPCR-MAPS: <u>m</u>ultiplexed <u>a</u>ssay with <u>p</u>athway <u>s</u>pecificity

To allow quantification of multi-modal receptor phenotypes for thousands of receptor variants in parallel, including full dose-response curves, we developed a modular platform that we call GPCR-MAPS. Our goal is to read out receptor states as directly as possible. In contrast to other approaches^27–30^, we therefore focused on quantifying direct binding interactions with receptors. Quantifying direct binding has proven powerful for understanding signaling by soluble proteins^35^ and GPCRs^36^ and avoids the confounding influence of non-linearities, feedback, and differential signal amplification when quantifying activity downstream of endogenous signaling networks.

GPCR-MAPS is compatible with cassette-based^37,38^ or SUNi mutagenesis^39^ to generate a single pooled library of receptor variants containing all possible single amino acid substitutions. The library is barcoded and rapidly integrated into a single locus in engineered HEK293 “landing pad” cells^40^ so that each cell expresses a single copy of a single variant.

In a first selection experiment, the cell surface expression of each variant is measured using FACS followed by sequencing^21,22,25^ to precisely quantify changes in receptor fold stability and trafficking (Figure 1a, Materials and Methods).

**Figure 1.**
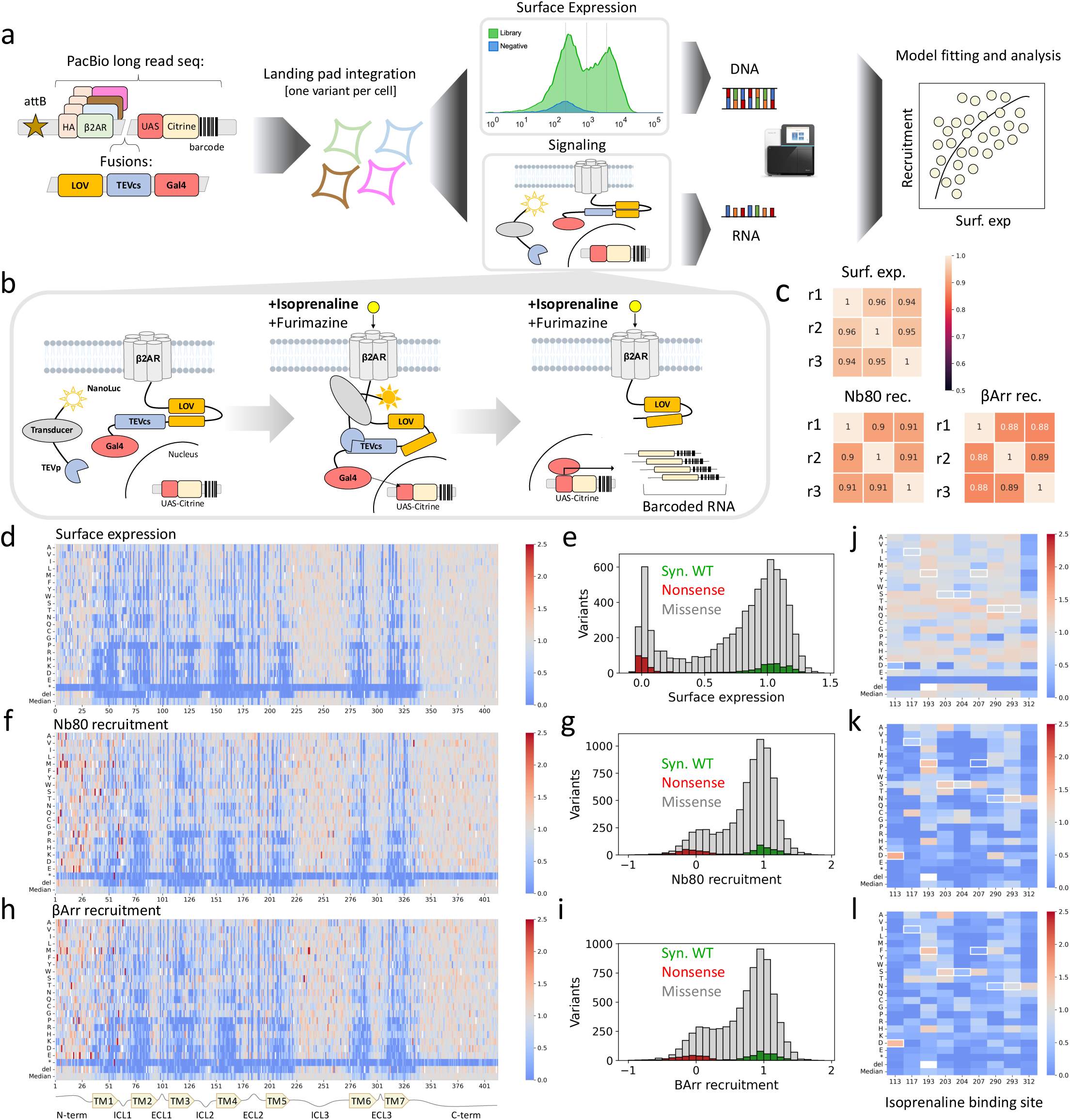
GPCR-MAPs. **a** Schematic of the experimental framework. A mutagenized GPCR (in this case β2AR) is encoded in a plasmid that also includes the attB recombination site, an N-terminal HA epitope tag, C-terminal fusions of a light, oxygen, and voltage sensitive (LOV) domain, the cleavage site of TEV protease (TEVcs), and Gal4 transcription factor. Downstream, on the same plasmid, is the fluorescent protein Citrine, driven by Gal4-responsive UAS promoter, with a degenerate DNA barcode included in the 3’ untranslated region. This construct is recombined into HEK 293T LLP-iCasp9-Blast landing pad cells. Then, two assays are performed to measure the surface expression (FACS-based) and recruitment activity (transcriptional reporter) of variants *en masse*. The outputs of these experiments (DNA for FACS and RNA for transcriptional reporter) are sequenced with short read technology. Estimated surface expression and recruitment scores are then modeled to understand the contribution of each of these to the variant effects. **b** Detailed schematic of the recruitment assay. The receptor construct is expressed from landing pad cells while the transducer construct is transiently transfected. Addition of agonist (isoprenaline in this case) and furimazine results in recruitment of transducer to the receptor fusion and subsequent cleavage of the TEVcs, liberating Gal4 which moves to the nucleus and drives transcription of the UAS-Citrine-barcode construct. **c** Replicate Pearson correlations for the surface expression, Nb80 recruitment, and βArr recruitment. p<10^-100^ in each case. **d** Heatmap representation of the surface expression data. β2AR positions on the x-axis, mutation on the y-axis. Gray cells indicate expression similar to wild-type while blue and red cells indicate reduced and increased expression, respectively. **e** Histogram of surface expression variant scores; nonsense variants in red, synonymous wild-type in green, and missense in grey. **f** Heatmap representation of the Nb80 recruitment data. Axes and color scale as in (d). **g** Histogram of Nb80 recruitment scores; colors as in (e). **h** Heatmap representation of β-Arrestin recruitment scores. Axes and color scale as in (d). **i** Histogram of β-Arrestin recruitment scores; colors as in (e). **j** Surface expression scores for residues in the isoprenaline binding site in the solved structure (PDB ID: 7DHR). **k** Nb80 recruitment scores for residues in the isoprenaline binding site. **l** β-Arrestin recruitment scores for residues in the isoprenaline binding site.

Additional selections then quantify direct binding of receptor interaction partners. To quantify recruitment of receptor interaction partners in response to ligand treatment we developed a modular protein interaction assay based on a previously described platform^41^ but extensively re-engineered to enable massively multiplexed experiments (see Materials and Methods). We also improved the signal to noise of the platform and extended it to allow modular read out of the recruitment of diverse GPCR protein interaction partners including the active-state-specific nanobody Nb80^42^, β-arrestin and miniG_s_^43^ (Supplementary Figure 1b).

The assay uses three genetic constructs to report on GPCR activation: 1) a GPCR of interest with a fusion of the exogenous transcription factor Gal4, linked by sequence encoding an evolved light, oxygen, and voltage sensitive (eLOV) domain as well as tobacco etch virus protease cleavage site (TEVcs), 2) a GPCR binding partner (e.g. β-arrestin) fused to NanoLuciferase (NLuc) and tobacco etch virus protease (TEV), and 3) the fluorescent protein Citrine under the control of the UAS promoter (Figure 1b and Supplementary Figure 1e). When co-expressed in cells, agonist-mediated activation of the GPCR causes recruitment of the binding partner to the receptor. In the absence of light, the eLOV domain cages the TEVcs and prevents cleavage; however, co-addition of the NLuc substrate furimazine results in bioluminescence, and when the transducer brings NLuc into close proximity with the eLOV domain, the steric hindrance is relieved and TEVp can cleave at TEVcs. This cleavage results in liberation of Gal4, which can move to the nucleus and drive transcription of the UAS-Citrine reporter construct, which is read out by short-read RNA sequencing (Figure 1b).

The recruitment assay allows the binding of multiple GCPR protein interaction partners to be quantified in a single condition (for example ligand concentration). However, because of its highly multiplexed RNA-seq based readout, it also allows the quantification of protein recruitment across many different conditions, for example ligand combinations, changes in culture conditions or, as we demonstrate here, ligand titrations allowing the construction of full dose-response curves for thousands of different receptors in a single pooled experiment.

### Application to the beta-2 adrenergic receptor (β2AR)

We generated a saturation mutagenesis library of β2AR in a vector also containing: an N-terminal HA epitope tag and C-terminal fusions of eLOV, TEVcs, and Gal4, and additionally the UAS-Citrine module (attB-HA-b2AR-eLOV-TEVcs-Gal4-UAS-Citrine, Supplementary Figure 1e) including all missense and nonsense substitutions, synonymous mutations where possible (i.e. the wild-type amino acid can be encoded by alternate codons), and single residue deletions. We then introduced short degenerate nucleotide barcodes into the 3’ UTR of Citrine and used long read sequencing to make 212,840 β2AR genotype-barcode links (Supplementary Figure 1c, Materials and Methods). These represented 8,970 out of an intended 9,046 unique genotypes (99.2%). We introduced this library into HEK293T landing pad cells^40^, ensuring a single variant is expressed per cell.

### β2AR surface expression

The first selection experiment of GPCR-MAPS is a FACS-based multiplexed surface expression experiment. Cells were immunostained with anti-HA antibody, then sorted into four bins based on fluorescence intensity (Supplementary Figure 1d, Materials and Methods). DNA from the cells sorted into each bin was purified and converted into sequencing libraries, and the barcodes were sequenced with short-read sequencing (Figure 1a, Materials and Methods).

All counts from barcodes representing the same variant were tallied and combined with the average fluorescence value of each FACS bin to arrive at surface expression estimates. Variants with low frequency in the library were filtered to yield 8,740 high confidence surface expression measurements that were highly reproducible across three replicates (Pearson r=0.94-0.96, Figure 1c, Materials and Methods, Supplementary Table 1). Scores were normalized between the median value of premature stop codons, set to 0, and the wild-type surface expression value, set to 1. A heatmap representation of the data shows nonsense mutations are uniformly damaging, except those occurring within intracellular loop 1 (ICL1) and those in the C-terminus, suggesting the receptor can be stably expressed as either a single-pass transmembrane protein, or as a C-terminal tail truncation. The transmembrane helices are more mutation sensitive than extra- or intra-cellular loops, consistent with previous work^21–25,28^ (Figure 1d). Synonymous wild-type and nonsense variants have well separated expression distributions, with missense variants spanning the full range between those two classes (Figure 1e). Our surface expression estimates are in line with previously estimated surface expression levels of alanine substitutions^18^ (Spearman’s ρ = 0.43, Supplementary Figure 1i).

### β-arrestin and Nb80 recruitment

The second selection experiments of GPCR-MAPS are protein interaction recruitment assays. For β2AR we quantified recruitment of two conformation-specific binders: Nb80 (as a surrogate for G-protein, as Nb80 returns higher signal to noise than miniG_s_ (Supplementary Figure 1b)) as well as β-arrestin. Namely, we transiently transfected one of the binder constructs (pUC19-NLuc-[Nb80/βArr]-uTEV1Δ, Supplementary Figure 1e) into cells bearing the β2AR receptor construct in the landing pad. We also included a condition where Gal4 was transfected alone (pUC19-CMV-Gal4, Supplementary Figure 1e); barcode frequencies in this control condition should be related to relative abundance in the library combined with any barcode-specific effects on RNA stability. The next day, saturating concentration of isoprenaline (10 μM) was added to the experimental samples to induce activation, and 24 hours later, those cells were collected. The Gal4 maximum induction samples were collected 30 hours after transfection. RNA was collected from all samples, reverse transcribed, converted to short-read sequencing libraries and sequenced (Supplementary Figure 1f-g, Materials and Methods).

Recruitment scores were calculated using the variant frequency in the experimental samples compared to variant frequency in the Gal4 condition and re-scaled with 0 representing the median of nonsense variants recruitment and 1 representing wild-type recruitment. Low frequency variants were filtered to yield 7,850 high confidence Nb80 and β-arrestin recruitment scores (Materials and Methods, Supplementary Table 1). Replicates were well correlated (Pearson’s r=0.9-0.91 for Nb80 replicates, r=0.88-0.89 for β-arrestin replicates, Figure 1c). Heatmap representations of the data reveal similar overall trends between Nb80 and β-arrestin (as well as with the surface expression data) defined by transmembrane regions being most sensitive to mutation (Figure 1f-h). Synonymous wild-type variants are again well separated from nonsense variants, and missense variants again span the whole distribution from nonsense to wild-type-like (Figure 1g, i). The Nb80 and β-arrestin recruitment scores are also correlated with each other (Spearman’s ρ = 0.87), as well as with previous multiplexed measurements of β2AR variants signaling through the G_s_ pathway^27^ (Spearman’s ρ = 0.44 and 0.47 for Nb80 and β-arrestin recruitment, respectively, Supplementary Figure 1j-k).

### Deconvolving recruitment and expression

Plotting Nb80 and β-arrestin recruitment scores against receptor expression reveals that changes in expression are an important cause of changes in recruitment (ρ=0.55, 0.6, for Nb80 and β-arrestin respectively, Figure 2a-b, Supplementary Figure 1h). However, some mutations have much larger effects on recruitment than can be explained by changes in expression. For example, missense mutations at the nine positions making up the isoprenaline binding site^44^ have minimal effect on expression (Figure 1j), while these same mutations are strongly deleterious for recruitment of both binding partners (Figure 1k-l).

**Figure 2.**
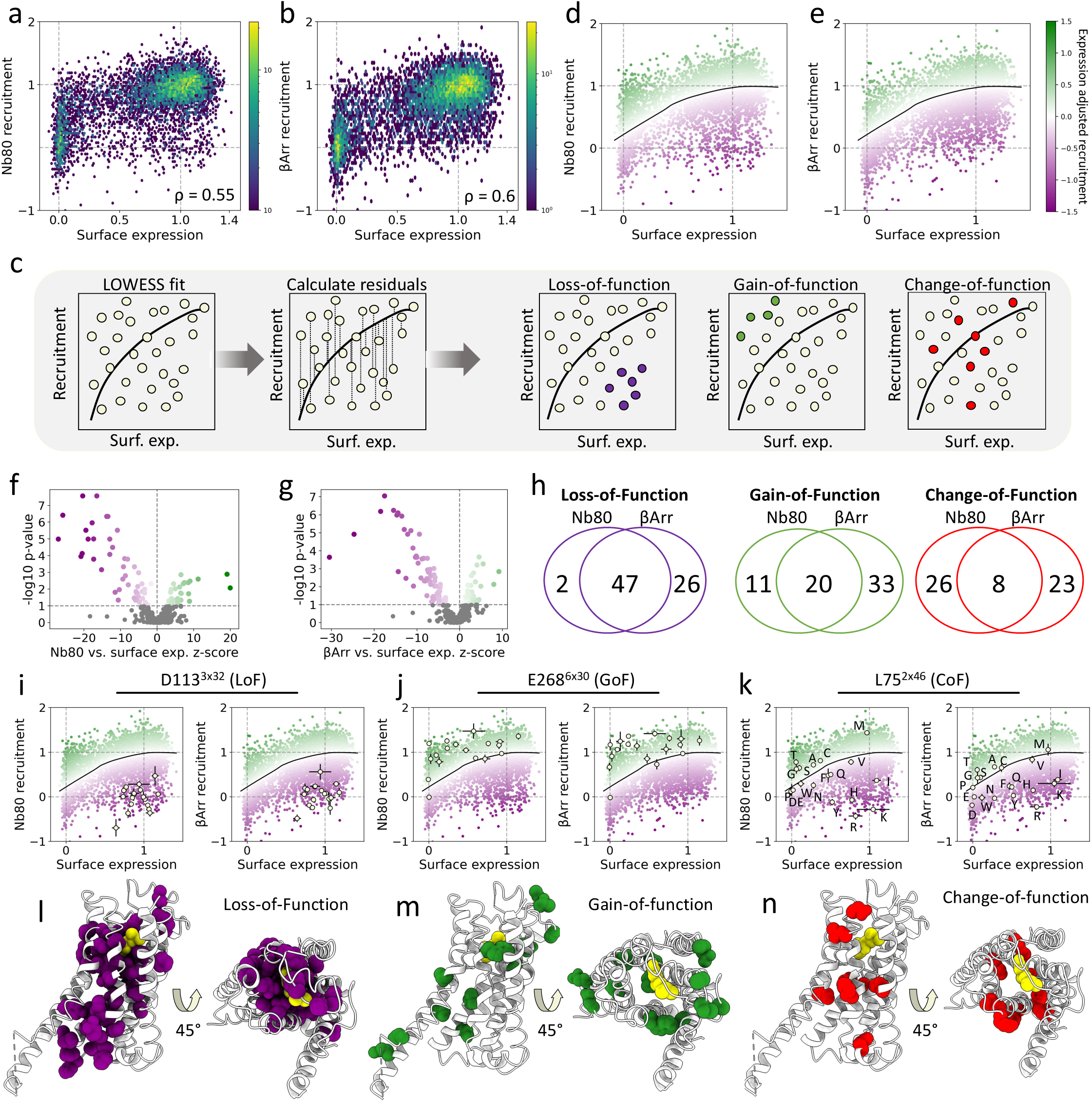
Deconvolving recruitment and expression. **a** Surface expression versus Nb80 recruitment scores for missense variants. p<10^-100^. **b** Surface expression versus β-Arrestin recruitment scores for missense variants. p<10^-100^. **c** Schematic of the method for identifying residues contributing to recruitment beyond what can be explained by surface expression. First, the recruitment data is related to surface expression with a LOWESS regression. Then, for each variant, residuals to this fit are calculated. Then, residues with significantly biased residuals are identified as loss-of-function or gain-of-function via Mann Whitney U test. Finally, positions where variants have a wide range of effects, quantified using the interquartile range of variant z-scores, are identified as change-of-function. **d** LOWESS fit of surface expression versus Nb80 recruitment, with variant points colored by residual to the fit. **d** Same as (e) but for β-Arrestin recruitment. **f** Volcano plot showing the position-average z-score and the minus log10 transformed p-value of all residues for Nb80 recruitment. Residues with significantly biased residuals are colored according to the average z-score. **g** Same as in (f) but for β-Arrestin recruitment. **h** Venn diagrams showing the number of residues called loss-, gain-, and chage-of-function for Nb80 and β-Arrestin recruitment. **i** Surface expression and recruitment effects of variants at D113^3x32^, plotted over the population of variant effects, for Nb80 and β-Arrestin recruitment versus surface expression. Error bars represent standard error of the mean. **j** Same as (i) but with variants at position E268^6x30^. **k** Same as (i) and (j) but with variants at position L75^2x46^. **l** Loss-of-function residues highlighted in purple on the isoprenaline-bound β2AR structure (PDB ID: 7DHR, isoprenaline in yellow). **m** Gain-of-function residues highlighted in green on β2AR. **n** Change-of-function residues highlighted in red on β2AR

### Residues contributing to recruitment beyond expression changes

The relationship between recruitment and expression is non-linear (Figure 2a-b). We therefore fit a LOWESS (locally weighted scatterplot smoothing) regression to the data, which acts as a null expectation of recruitment produced by a given surface expression level. This null expectation explains a significant amount of variance (R^2^ = 0.37 and 0.42 for Nb80 and β-arrestin, respectively).

We next quantified changes in recruitment not explained by expression changes by calculating recruitment residuals and subsequently z-scores of all variants to the global model (i.e. residual divided by standard error of the mean for each variant, Figure 2c-e). Comparing the group of z-scores at each position with the z-scores from all other positions with a two-sided Mann-Whitney U test allows us to identify residues where mutations have greater or lesser recruitment than would be expected based on their surface expression level (Supplementary Table 2).

### Complete functional maps of Nb80 and β-arrestin recruitment

For both Nb80 and β-arrestin, more residues confer loss-of-function (LoF) than gain-of-function (GoF) when mutated, and the magnitude of loss is greater than the magnitude of gain (Figure 2f-g). In total, 49 and 73 residues are LoF for Nb80 and β-arrestin, respectively (false discovery rate, FDR = 0.1, Supplementary Table 2). Of these, 47 are shared (Figure 2h, OR=304.6, p=2.2x10^-40^, two-sided Fisher exact test). An example of a strong LoF residue is D113^3x32^ (Figure 2i, GPCRdb numbering^45^ in superscript), which participates directly in isoprenaline binding. There are 31 and 53 GoF residues for Nb80 and β-arrestin (FDR=0.1), respectively, of which 20 are shared (OR=19.2, p=1.3x10^-12^, two-sided Fisher exact test). E268^6x30^ is the most significant GoF (Figure 2j), consistent with previous work demonstrating constitutively activating mutations at this site^46^. Finally, we extended this analysis to highlight residues where mutations have highly variable effects on recruitment. Namely, after excluding the LoF and GoF residues, we compared the interquartile range (IQR) of z-scores at all positions (Supplementary Figure 2a-b). We categorized those residues with IQR above the 90^th^ percentile as change-of-function (CoF): 34 and 31 residues for Nb80 and β-arrestin, respectively, with 8 shared (Figure 2h, OR=2.9, p=0.037). An example of a strong CoF residue is L75^2x46^ (Figure 2k), which is near the allosteric sodium binding site and neighbors LoF residue A76^2x47^. At this position, mutations to small, uncharged residues (G, S, T, A, C) decrease surface expression, but increase Nb80 and β-arrestin recruitment relative to the modeled expectation. On the other hand, positively charged residues (H, R, K) are well expressed but strongly decrease recruitment of both proteins.

Average position z-scores and position IQRs are well correlated (r = 0.73 and 0.56, respectively, Supplementary Figure 2c-d), indicating that many residues contribute similarly to recruitment of both proteins. Plotting the residues identified for both proteins onto the structure of isoprenaline-stimulated β2AR (PDB ID: 7DHR) reveals that LoF residues are predominantly within the core of the receptor (Figure 2l), whereas GoF and CoF residues are dispersed on the periphery (Figure 2m-n).

### Loss-of-function residues define a core activation network

The majority of the shared LoF residues (39 out of 47) form an interconnected path linking the ligand binding site all the way to the transducer binding site in the active state receptor structure (Figure 3a, hereafter referred to as the “core activation network”). All 9 residues of the isoprenaline binding pocket are identified (D113^3x32^, V117^3x36^, F193^45x52^, S203^5x43^, S204^5x44^, S207^5x461^, F290^6x52^, N293^6x55^, and N312^7x38^) along with many residues in the second shell of the ligand binding pocket (i.e. residues that contact residues of the isoprenaline binding site, Figure 3b). These residues include W286^6x48^ of the CWxP motif and I121^3x40^ of the PIF motif^47,48^. The core activation network then connects to N322^7x49^ of the critical NPxxY motif^47^ via two pathways: first, W286^6x48^ contacts N318^7x45^ which then connects to N322^7x49^. Second, a pathway from A119^3x38^ goes through A78^2x49^ and D79^2x50^ of the allosteric sodium binding pocket which then contacts N322^7x49^. Of the NPxxY motif, N322^7x49^, I325^7x52^, Y326^7x53^ are identified as LoF. Finally, in the active state structure, Y326^7x53^ of the NPxxY motif then contacts R131^3x50^ (of the DRY motif) and I127^3x46^ of helix 3. Y326^7x53^ is also thought to make an important water-mediated hydrogen bond with Y219^5.58^,^49^ which is also identified as LoF and makes a direct contact with R131^3x50^.

**Figure 3.**
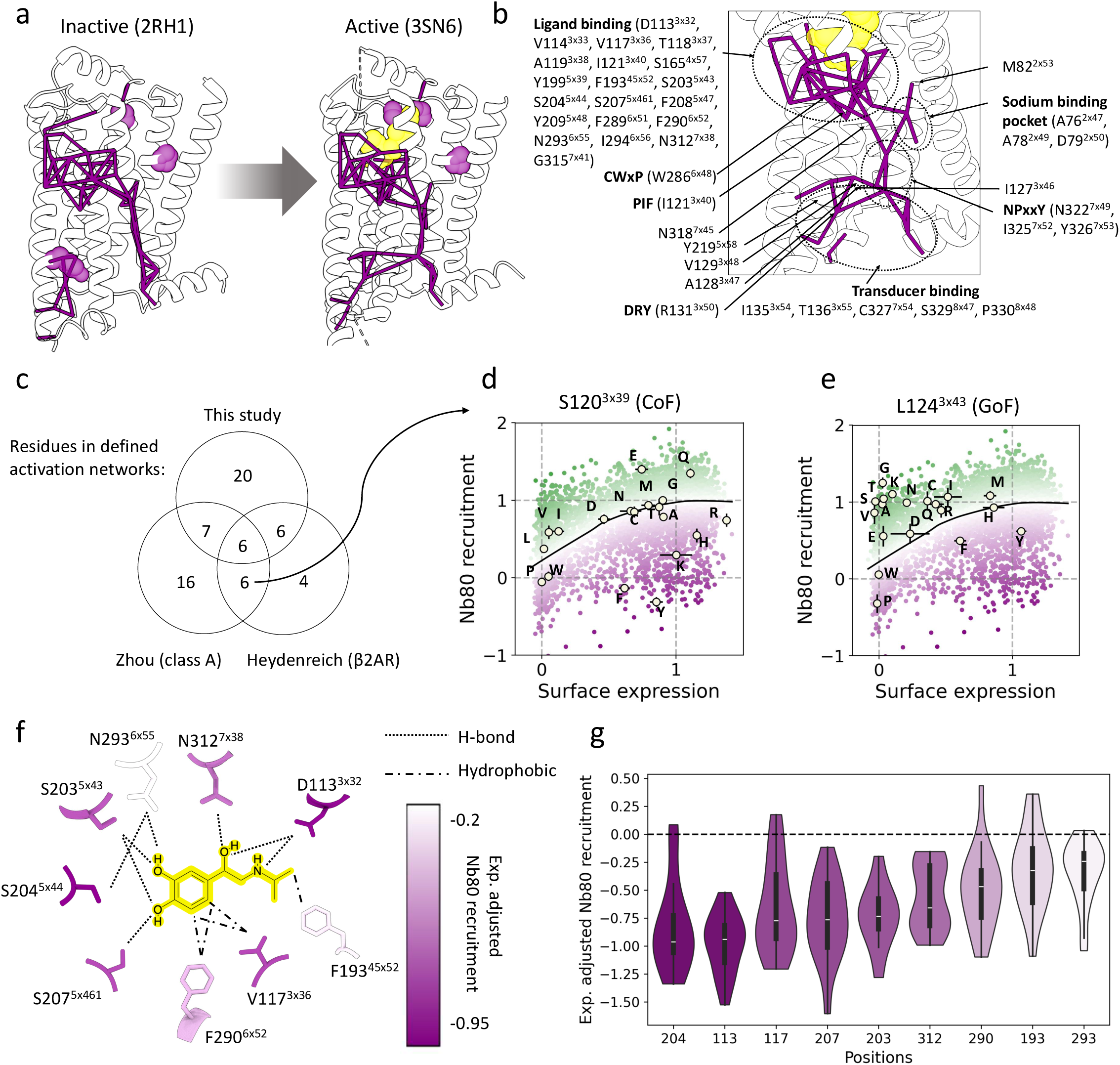
A core activation network links ligands to transducers. **a** Inactive and active state structures of β2AR with LoF residues colored purple. Those LoF residues that make a direct contact with other LoF residues are visualized as pseudobonds connecting them, while disconnected LoF residues are shown with spheres. **b** Enlarged view of the core activation network with residues labeled and their participation in different structural motifs. **c** Venn diagram showing the overlap between network residues identified in this work and previous work. **d** Variant surface expression and Nb80 recruitment at S120^3x39^, a CoF residue, are plotted against the population of all variants. **e** Same as (d) but with variants at L124^3x43^, a GoF residue. **f** The isoprenaline binding pocket, with isoprenaline highlighted in yellow, and the residues making up the binding site colored according to their median expression adjusted Nb80 recruitment level. **g** Violin plot of expression adjusted Nb80 recruitment at the isoprenaline binding site residues. Violins are colored according to their median value, with the same color scale as in (f).

Of the eight LoF residues not connected to the core activation network, three likely directly disrupt transducer binding^42,50^ (P138^34x50^, F139^34x51^, and Y141^34x53^), while three more take part in critical disulfide bonds between ECL2 and TM3 (C106^3x25^, C190^ECL2^, C191^45x50^). F193^45x52^ is a disconnected LoF residue that makes direct contact with bound isoprenaline. Finally, P88^2x58^ in TM2 is the last disconnected residue.

Previous work has used structural analysis to define an activation network for class A GPCRs^51^ and alanine scanning mutagenesis to define an activation network for β2AR^18^. About half of the residues identified in our network are present in at least one of the others (19/39, Figure 3c). We identify more residues near the ligand binding site compared to the other networks (Supplementary Figure 2e-g). Interestingly, of the six residues shared by the other two networks that are not part of our core activation network, we identify four of these as either CoF (S120^3x39^ and F332^8x50^) or GoF (L124^3x43^ and D130^3x43^), emphasizing the value of empirically assaying all 19 substitutions per position. At S120^3x39^, which neighbors I121^3x40^ of the PIF motif, positively charged substitutions (R, K, H) yield stable receptors with significantly reduced recruitment, while the E and Q also yield stable receptors but significantly increase recruitment (Figure 3d). At L124^3x43^, most variants decrease surface expression but increase relative recruitment, with the exception of the aromatic amino acids (W, F, and Y) and proline which recruit Nb80 less than expected given their expression level (Figure 3e).

### Gain-of-function sites are located at the ionic lock, near the NpxxY motif, and in the extracellular vestibule

We next examined the location of the GoF residues (Supplementary Figure 3a, f). Two of the GoF residues, D130^3x49^ and E268^6x30^, participate in the highly conserved ionic lock (with R131^3x50^ of the DRY motif^52^). While R131^3x50^ of the DRY motif is a LoF residue that contacts Nb80^42^ and is predicted to contact β-arrestin,^50^ some mutations at D130^3x49^ of the DRY motif have been shown to constitutively activate the receptor^53^. The other GoF residues are predominantly in two clusters: one consisting of TMs 3, 5, and 6, near the NPxxY and DRY motifs (Supplementary Figure 3b). L124 sits in the center of the receptor, near the I of PIF motif, DRY motif, and NPxxY. A few mutations at L124^3x43^ have been shown to cause constitutive activity at this site^54^ but we show that most variants activate more than expected (discussed above). The other cluster is near the ligand binding site and in the extracellular vestibule (Supplementary Figure 3c). E187 and T189 are in ECL2, where they may affect ligand binding or activation^55^. W313^7x39^ and V87^2x57^ form two van der Waals interactions in the inactive (PDB:2RH1), but only one in the active state (PDB:3SN6, GetContacts); our data suggest that disrupting this interaction leads to increased recruitment. Finally, there are 7 GoF residues in the C-terminal tail (Supplementary Figure 3g).

### Change-of-function sites are adjacent to the core network

Plotting the CoF residues onto the inactive and active state structures (Supplementary Figure 3d) reveals that they cluster near important LoF residues that make structural changes during activation: for instance, S120^3x39^ and E122^3x41^ are adjacent to I121^3x40^ of the dynamic PIF motif. Likewise, F321^7x48^ is adjacent to N322^7x49^ of the dynamic NPxxY motif and identified LoF, while F332^8x50^ is nearby (Supplementary Figure 3e). This proximity to essential switch residues suggests that CoF residues may specifically stabilize or destabilize the active or inactive receptor state depending on the amino acid substitution.

### Differential contribution of residues in the isoprenaline binding pocket

Next, we examined the effects of substitutions at residues making up the isoprenaline binding site. While all residues are LoF, the magnitude of effects differs among the constituents. D113^3x32^, which forms two hydrogen bonds with isoprenaline, has strongly damaging effects (median expression-adjusted Nb80 and β-arrestin recruitment = -0.94 and -0.89, Figure 3f-g and Supplementary Figure 3h-i). N293^6x55^ also forms a hydrogen bond with isoprenaline, but substitutions here are the least damaging of the binding site (median expression-adjusted Nb80 and β-arrestin recruitment = -0.24 and -0.4). This tolerance is surprising as N293^6x55^ also forms a hydrogen bond with S204^5x44^, which doesn’t directly contact the ligand but has strongly deleterious effects (median expression-adjusted Nb80 and β-arrestin recruitment = -0.96 and -0.81). This suggests the role of S204^5x44^ may be more related to receptor activation than ligand binding.

### Genetic control of receptor recruitment bias

Next, we directly compared mutational effects on Nb80 and β-arrestin recruitment (Figure 4a). Missense mutational effects on recruitment of Nb80 and β-arrestin are well correlated (Pearson’s r=0.89, Figure 4b). However, single mutants previously shown to bias the receptor toward G protein signaling (F193A^56^, Y219A^57^, and Y199A^58^) are indeed biased towards Nb80 in our experiments (Figure 4c). To find residues with stronger effects on one pathway compared to the other, we again fitted a LOWESS regression curve to the data, and calculated residuals and subsequently z-scores for all variants in the dataset. Then, we compared z-scores of mutations at each position with z-scores of mutations at all other positions. This identifies 19 residues where mutations have significantly greater effects on β-arrestin recruitment and 7 positions where mutations have significantly greater effects on Nb80 recruitment (FDR=0.1, Figure 4d, Supplementary Table 3). As before, we then compared the IQR of all remaining residues; here, to more closely match the number of biased residues, we prioritized the top 5% of residues as recruitment bias switches (Supplementary Figure 4a).

**Figure 4.**
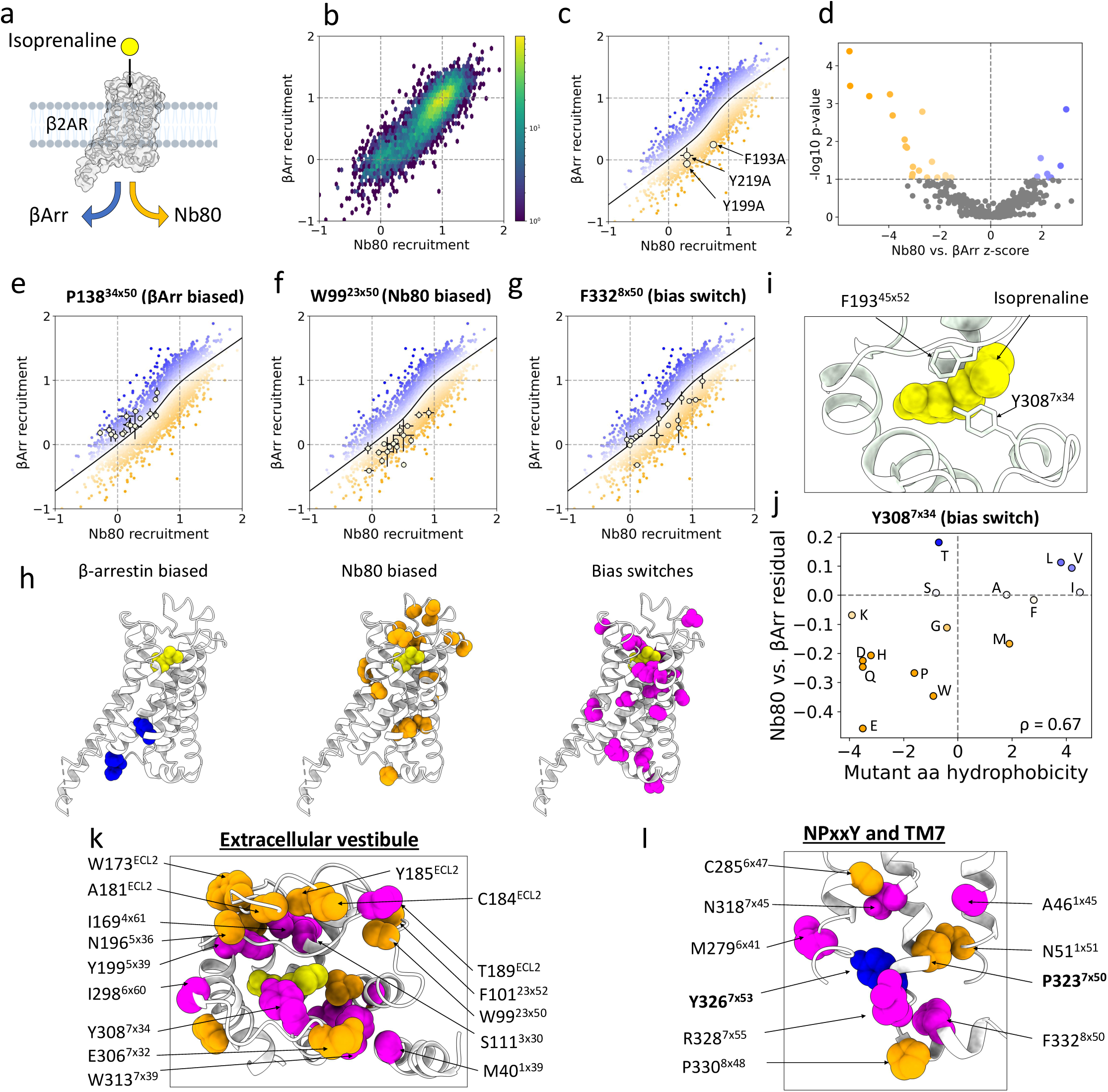
Genetic control of receptor recruitment bias. **a** Schematic showing β2AR activation through β-Arrestin and Nb80 in response to isoprenaline. **b** Nb80 versus β-Arrestin recruitment for missense variants. p<10^-100^, n=6,791. **c** Nb80 versus β-Arrestin recruitment with a LOWESS fit and variant scores colored by residual to the fit line. Individual variant scores, with error bars representing standard error of the mean, shown for three variants previously reported to cause biased signaling. **d** Volcano plot showing the position-average z-score and the minus log10 transformed p-value of all residues for Nb80 versus β-Arrestin recruitment. Residues with significantly biased residuals are colored according to the average z-score. **e** Nb80 and β-Arrestin recruitment effects of variants at W99^23x50^, plotted over the population of variant effects, error bars represent standard error of the mean. **f** Same as in (e) but with variants at P138^34x50^. **g** Same as in (e) but with variants at F332^8x50^. **h** β-Arrestin biased, Nb80 biased, or bias switch residues highlighted on the solved structure, 7DHR. Isoprenaline shown in yellow. **i** F193^45x52^ and Y318^7x34^ form a “lid” over isoprenaline in the solved structure (7DHR). **j** Hydrophobicity (Kyte-Doolittle scale) of the mutant amino acid compared with the Nb80 versus β-Arrestin recruitment bias. **k** Enlarged view of the extracellular vestibule. Colored residues are significantly Nb80-biased (orange), β-Arrestin-biased (blue), or bias switches (magenta). **l** Enlarged view of NPxxY motif and TM7, colored residues same as in (k).

Examples of strongly biased residues are shown (Figure 4e-f) as well as a strong bias switching residue (Figure 4g). One expectation is that mutations at residues making direct contacts with either Nb80 or β-arrestin could result in biased recruitment. Nb80 and β-arrestin have overlapping but distinct binding interactions with the intracellular portion of β2AR. Nb80 forms direct contacts with 16 residues in TMs 3, 5, 6, and 7^42^. Since there is no experimentally determined structure of β2AR bound by β-arrestin, we defined this interface from a simulation study^50^. The β-arrestin binding interface is predicted to be larger, encompassing 28 residues across ICLs 1, 2, 3, and TMs 6 and 7, with 10 residues overlapping the Nb80 interface.

Three out of the seven residues biased toward β-arrestin recruitment make direct binding interactions with both Nb80 in the solved structure and with β-arrestin in the simulation (P138^34x50^, F139^34x51^, and Y326^7x53^), suggesting a more critical role for these residues in binding Nb80. The other four β-arrestin-biased residues are either in ICL3 (G252 and G255) or the C-terminal tail (E373 and C406, Supplementary Figure 4b). The only residue biased toward Nb80 within either binding interface is P330^8x48^, which is predicted to interact with Nb80 but not β-arrestin. There is also one bias switch residue, R328^7x55^, that is predicted to only bind β-arrestin. This relative sparsity of bias effects suggests that few residues at the transducer interface contribute significantly more to recruitment of each protein (Supplementary Figure 4c).

### The extracellular vestibule allosterically controls recruitment bias

In contrast, there are many residues biased toward Nb80 recruitment located away from the orthosteric site in the extracellular vestibule: W99^23x50^, F101^23x52^, G102^3x21^, W173^ECL2^, A181^ECL2^, C184 ^ECL2^, Y185 ^ECL2^, N196^5x36^, and E306^7x32^ (Figure 4h, k). The most strongly Nb80 biased residue, W99^23x50^, was previously identified as particularly mutationally intolerant^27^; however only by measuring recruitment of both proteins could we uncover its strongly biased effects (Figure 4f). Considered together, the extracellular loops (ECLs) are enriched in Nb80 biased residues compared to the whole receptor (OR=6.4, p=1.0x10^-3^, one-sided Fisher exact test), and in particular, ECL2 is enriched for Nb80 biased residues (OR=5.6, p=0.014, one-sided Fisher exact test).

The importance of the extracellular vestibule for G-protein biased activation revealed in our mutational data is also consistent with structural data. Salmeterol is a G-protein biased β2AR agonist that binds the orthosteric site in a similar way as the endogenous ligand (epinephrine) except that it has a long aryloxyalkyl tail that extends into the extracellular vestibule, where it interacts extensively with ECL2^59^. Also, comparative structural analyses of other GPCRs have suggested that the extracellular vestibule adopts a more constricted conformation in the β-arrestin-bound state compared to the G-protein-bound state.^60^ Taken together, these data point to the extracellular vestibule, and in particular ECL2, as hotspots for G-protein-biased agonism at the β2AR.

Among the bias switching residues, the second strongest is the vestibule residue, Y308^7x34^ (Supplementary Figure 4d). This residue has been proposed to be necessary for G-protein signaling^61^, as well as forming a hydrophobic gate with F193^45x52^ that interacts with and dehydrates ligand molecules as they traverse the vestibule and approach the orthosteric binding site^62^. In the active state, these two residues form a “lid” over the ligand, with F193^45x52^ making a direct contact with isoprenaline (Figure 4i). Based on the alanine mutation, F193^45x52^ was postulated to coordinate β-arrestin signaling^56^. Interestingly, while this single variant does have a stronger effect on β-arrestin recruitment, other variants at the same site have the opposite effect (Supplementary Figure 4e). We examined the effects of mutations at these two sites and found that recruitment bias is correlated with mutant hydrophobicity for Y308^7x34^ (ρ = 0.67, p = 4.3x10^-3^, Figure 4j), but not for F193^45x52^ (ρ = -0.36, p = 0.16, Supplementary Figure 4f). This suggests that the role of Y308^7x34^ in determining bias is directly tied to its hydrophobicity, while the relationship of F193^45x52^ to bias is more nuanced.

### Recruitment bias in TM7

Besides the vestibule, there are multiple bias-functional residues (Nb80- or β-arrestin-biased or bias switches) in or near the NPxxY motif of TM7(Figure 4l). In fact, TM7 is enriched for bias-functional residues compared to the whole receptor (OR=3.7, p=0.011, one-sided Fisher exact test, Supplementary Figure 4g). P323^7x50^ is not predicted to contact directly β-arrestin^50^, suggesting that the Nb80 bias of the residue cannot be explained simply by disrupting a direct contact. N318^7x45^, F332^8x50^, and R328^7x55^ are all bias switches in close proximity to the NPxxY motif (Figure 4k). A role for TM7 in determining bias is consistent with prior nuclear magnetic resonance experiments demonstrating changes in TM7 conformation upon binding of biased agonists^63^.

### Activation functions for >7,500 β2AR variants

The data presented thus far quantifies recruitment to β2AR with saturating levels of isoprenaline, which convolutes the contribution of different pharmacological parameters, such as potency (the amount of agonist needed to elicit a certain response) and efficacy (the maximum recruitment response elicited by saturating concentrations of agonist). However, because of its pooled design and RNA-seq based output, GPCR-MAPS can also be used to quantify the full dose-response curves (activation functions) of all receptor variants in a library.

To demonstrate this capability, we performed the Nb80 GPCR-MAPS recruitment assay with 12 serially diluted concentrations of isoprenaline, ranging from 10 pM to 10 uM (Figure 5a). In order to compare the magnitude of recruitment across experimental conditions, following RNA isolation, we performed quantitative PCR (qPCR) to estimate the total amount of reporter transcription (Figure 5b, Supplementary Figure 5a, Materials and Methods). Then, by deep sequencing the barcodes and normalizing the frequencies by the estimated total reporter RNA, we arrived at normalized scores across the 12 concentrations (Figure 5c and Supplementary Figure 5b, Materials and Methods). This allowed us to quantify dose-response relationships for 7,680 different receptor variants.

**Figure 5.**
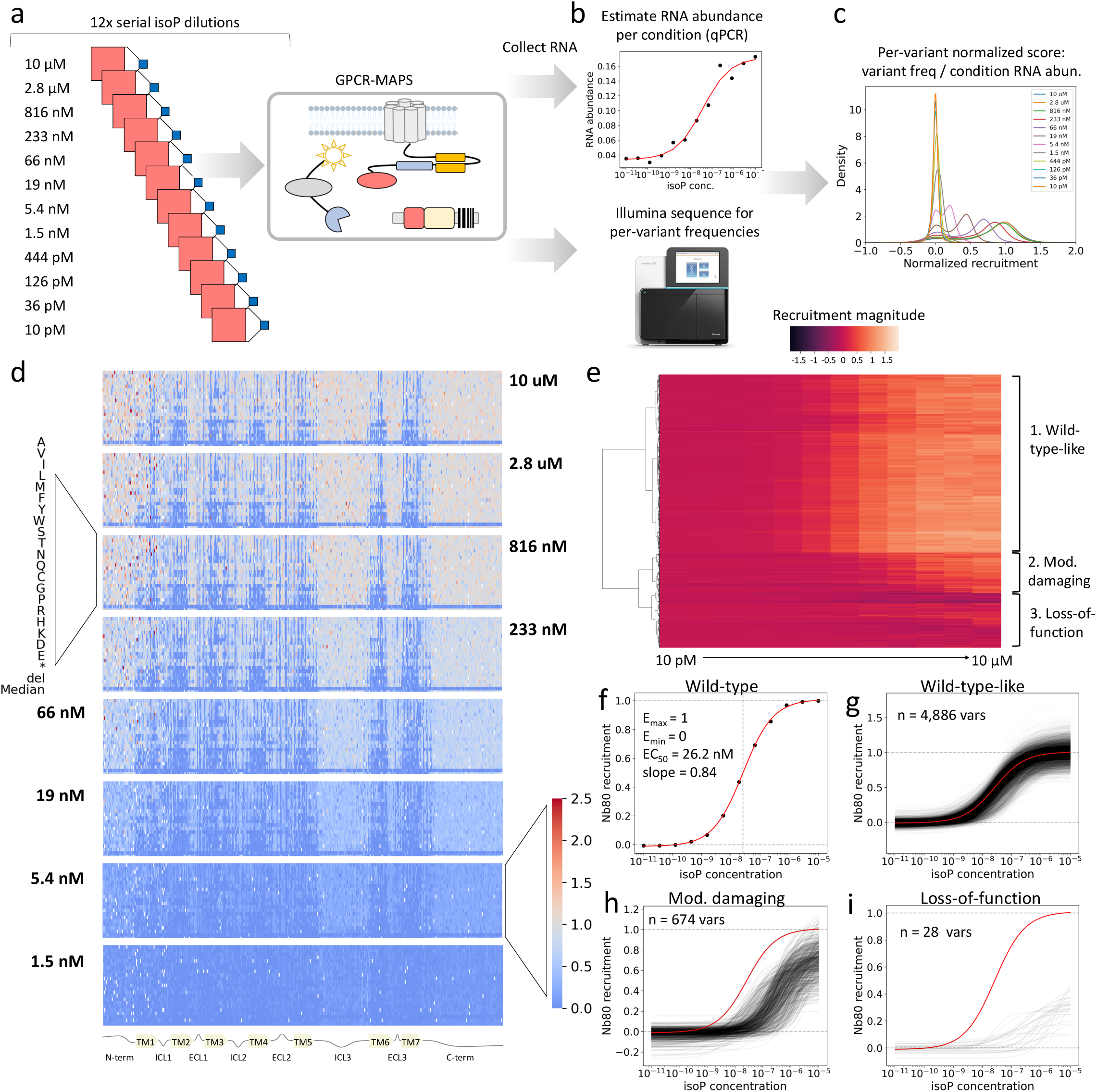
Quantifying >7,500 dose-response relationships. **a** Experimental design: serial dilutions of isoprenaline were used to stimulate landing pad cells expressing the β2AR library. Then, the GPCR-MAPS recruitment assay was performed and RNA was collected from the stimulated cells. **b** The RNA abundance from each sample was first quantified with qPCR, then the variant frequencies were estimated with short-read sequencing. **c** Per-variant normalized scores were then calculated by normalizing variant frequencies in each condition by the RNA abundance of each condition. **d** Heatmap representations of the 8 highest concentration drug treatment conditions. β2AR positions on the x-axis, mutation identities on the y-axis. One y-axis is blown up; the rest of the heatmaps follow this same scheme. Grey cells indicate wild-type-like recruitment, blue and red indicate reduced and increased recruitment, respectively. One color bar is blown up; the rest of the heatmaps follow this same scheme. **e** Clustered heatmap of recruitment intensities. Color bar on top. Three main clusters correspond to wild-type-like, moderately damaging, and complete loss-of-function curves. **f** The curve fit to the wild-type data, with E_max_, E_min_, EC_50_, and slope indicated. **g** Fit curves from the wild-type-like cluster in black, wild-type curve in red. **h** Fit curves from the moderately damaging cluster in black, wild-type curve in red. **i** Fit curves from the loss-of-function cluster in black, wild-type curve in red.

Heatmap and correlation representations of the data reveals induction of Nb80 recruitment across the ∼92,000 measurements (Figure 5d and Supplementary Figure 5c-d). Clustering the induction dataset revealed three predominant clusters-these can be assigned as wild-type-like, moderately damaging, and strong loss-of-function (Figure 5e).

Next, we fit a four-parameter Hill curve to the wild-type genotype, normalizing E_min_ and E_max_ to 0 and 1, respectively, and deriving a half maximal response (EC_50_) estimate of 26.2 nM and slope (Hill coefficient) of 0.84 (Figure 5f). Then we fit curves to all variants; after quality control (Supplementary Figure 5e-f, Materials and Methods), this resulted in 5,588 high quality dose-response curves with well-estimated parameters (Figure 5g-i). We note that variants with reduced activation level are more poorly fit and therefore likelier to be filtered (Supplementary Figure 5e). Our EC_50_ measurements correlate well with the results of individually assayed substitutions^18^ (Pearson r=0.74, n=317 variants, Figure 6b).

**Figure 6.**
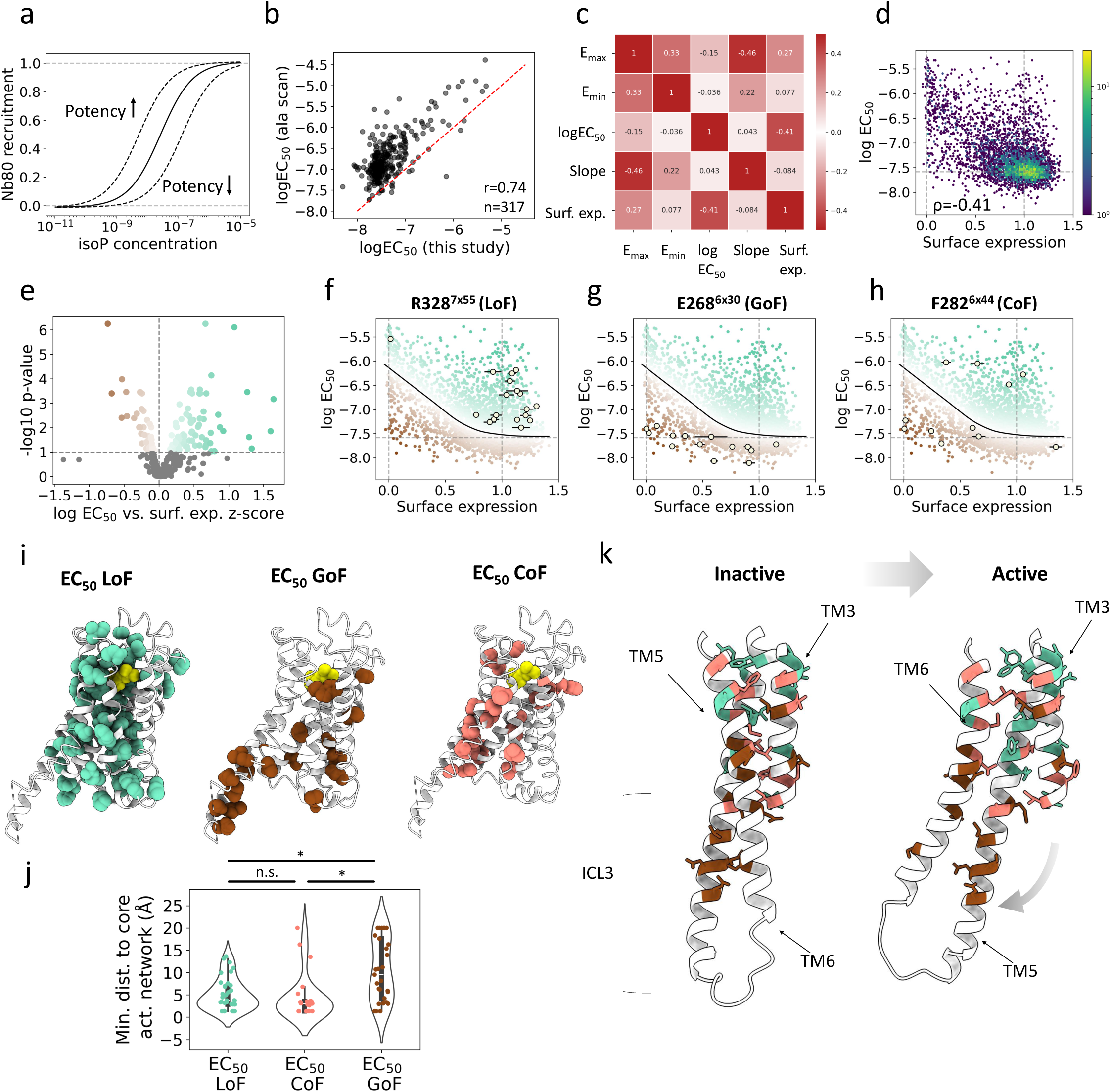
Genetic determinants of potency. **a** Schematic showing example curves for increased and decreased potency. **b** Comparison of variant EC_50_ estimates from this study with previously published estimates^18^. Red dashed line is x=y line. **c** Spearman correlations of curve fitting parameter estimates and surface expression scores. **d** Surface expression scores compared with log-transformed EC_50_ estimates. p<10^-100^. **e** Volcano plot showing the position-average z-score and the minus log10 transformed p-value of all residues for log EC_50_ versus surface expression. **f** Log-transformed EC_50_ versus surface expression for variants at R328^7x55^, plotted over the population of variant effects, error bars represent standard error of the mean. **g** Same as in (f) but with variants at E268^6x30^. **h** Same as in (f) but with variants at F282^6x44^. **i** EC_50_ loss-of-function, gain-of-function, or change-of-function residues highlighted on the solved structure, 7DHR. Isoprenaline show in yellow. **j** Violin plots showing minimum distance between LoF, GoF, or CoF residues from the core activation network in AlphaFold-Multistate active state model. Distances capped at 20Å. **k** Portions of TMs 3, 5, and 6, as predicted by AlphaFold-Multistate and downloaded from GPCRdb^45^. LoF residues colored aquamarine, GoF residues colored brown, and CoF residues colored salmon.

### Surface expression is a major determinant of receptor potency

Changes in potency are often measured by comparing EC_50_ values (Figure 6a). To investigate determinants of potency, we first quantified the correlations between the four Hill function parameters, as well as their dependence upon surface expression. Expression level has been predicted to be an important influence on signaling parameters by theoretical work^64–66^, and by experimental work on a plant hormone receptor^35^. The strongest relationships we found were between slope and E_max_ (ρ = -0.46) and between surface expression and log EC_50_ (ρ = -0.41, Figure 6c-d). Surface expression level is thus a major influence on potency in the β2AR. To mechanistically understand changes in potency it is therefore necessary to again quantitatively deconvolve direct mutational effects from those mediated by changes in receptor expression.

### Potency tuning

To isolate mutational effects on EC_50_ beyond those caused by changes in expression we again used LOWESS regression to model the relationship between surface expression and EC_50_ (Supplementary Figure 6a). We quantified residuals to that model to identify positions where mutations have consistently higher or lower EC_50_ than expected from their impact on expression. 29 residues were identified as GoF (i.e. they had EC_50_ lower than expected) and 59 were identified as LoF (i.e. higher EC_50_ than expected, FDR=0.1, Figure 6e-f). There is overlap between these sets and the sets identified from the saturating concentration experiment (23/59 LoF shared, 4/29 GoF shared). Finally, the interquartile range was used again to identify 29 EC_50_ CoF residues (one shared with saturating concentration experiment, Supplementary Figure 6b, Supplementary Table 4).

In principle, mutations could affect potency beyond changes in expression by changing the affinity of isoprenaline binding, or by affecting the probability of isoprenaline-bound receptor to adopt the active conformation^67^. While mutations at residues making direct contact with isoprenaline would be expected to reduce affinity, mutations at these sites mostly totally abolish activation; dose-response curves could only be fit to 32 out of 132 missense variants at these sites. However, these 32 variants do have over an order of magnitude reduced potency (median EC_50_=477 nM, compared to wild-type 26.2 nM).

Interestingly, EC_50_ LoF residues are enriched in the second shell (15/59 LoF sites (25.4%, OR=4.12, p=1.9x10^-4^, Fisher exact test). While not statistically enriched, a higher fraction of EC_50_ CoF sites (5/24, 20.8%) are in the second shell compared to only 1 out of 29 GoF residues (3.4%) in the second shell (Figure 6i). This shows that whereas mutations in residues directly contacting the ligand have very strong effects on affinity and so completely inactivate recruitment, mutations in second shell residues frequently can have more moderate effects and modulate the response. These effects are, by definition, allosteric, either indirectly altering ligand binding affinity or altering the consequences of ligand engagement. This also suggests that mutations in the second shell are likelier to reduce affinity than increase it.

Beyond changes in ligand affinity mutations can also affect recruitment by altering the likelihood of ligand-bound receptor adopting the active conformation^67^. One way to disrupt the activation pathway could be for mutations to interfere with the core activation network identified in the maximum recruitment experiment. We therefore quantified the distance of EC_50_-affecting residues to this network and found that LoF and CoF residues are closer in 3D space than GoF residues (p=7.4x10^-4^, 1.3x10^-3^, respectively, two-sided Mann Whitney U test, Figure 6j). Taken together, as with the maximum recruitment analysis, these data suggest that mutations that reduce potency do so by perturbing the core activation pathway whereas the mechanism-of-action of GoF mutations is likely distinct.

Three of the four most significant EC_50_ GoF residues are E268^6x^^30^ (Figure 6g), D130^3x49^, and L272^6x34^. E268^6x30^ and D130^3x49^ were already identified in the maximum recruitment analysis and make up the conserved ionic lock while L272^6x34^ is in close proximity to the ionic lock, where mutations are also thought to destabilize the inactive state^67^. Nearby E268^6x30^ in intracellular loop 3 (ICL3), there are nine EC_50_ GoF residues, which is enriched compared to the whole receptor (OR=3.38, p=7.3x10^-3^, one-sided Fisher exact test). This loop connects TM5 and TM6, which make dramatic movements upon receptor activation (Figure 6k). These GoF residues could act in a similar way as D130^3x49^, E268^6x30^, and L272^6x34^, weakening the interaction between TM3 and TM6 or other helices. Alternatively, previous studies suggest that ICL3 may have an autoinhibitory role that sterically occludes transducer binding in the inactive state, and that some mutations can abrogate this autoinhibitory effect^68^. Further experiments would be necessary to determine the exact mechanism of the EC_50_ GoF phenotypes in ICL3. Additionally, there are two GoF residues in the N-terminus that coincide with the two N-glycosylation sites^69^ (N5 and N15, Supplementary Figure 6d) suggesting that loss of these glycosylation modifications increases receptor potency.

F282^6x^^44^ of the PIF motif is the strongest EC_50_ CoF residue (Figure 6h) consistent with previous data showing multiple constitutive activating mutations and at least one attenuating mutation^70^. Likewise, S120^3x39^ and L212^5x51^ are identified as CoF and are directly adjacent to the other two members of the PIF motif (I121^3x40^ and P211^5x50^) reinforcing that some perturbations to this key switch motif can result in gains or loss of recruitment. There are also several EC_50_ CoF residues in TMs 5 and 6 (4 in each, Supplementary Figure 6c), reflecting the key role of these helices in activation.

### Annotation of surface pockets

Finally, we used our β2AR data to illustrate how GPCR-MAPS can be used to functionally annotate and prioritize potential pockets in receptors. AS408 is a negative allosteric modulator (NAM) that binds away from the orthosteric and transducer binding sites, between TMs 3 and 5^71^ (Figure 7a). Across the three analyses (maximum recruitment, biased recruitment, and EC_50_), the 7 residues making up this pocket have 7 total annotations, compared to 96 annotations for all 220 structurally resolved surface exposed residues (>10% relative solvent accessible surface area in 2RH1), indicating a functionally important pocket. In particular, V129^3x48^ is LoF for maximum recruitment as well as EC_50_, while E122^3x41^ is CoF for maximum recruitment as well as EC_50_. In addition, C125^3x44^ is CoF for EC_50_ and I214^5x53^ is LoF for EC_50_. Mutations at P211^5x50^ are biased toward Nb80 recruitment, though AS408 is not thought to be a biased NAM (Supplementary Figure 7a).

**Figure 7.**
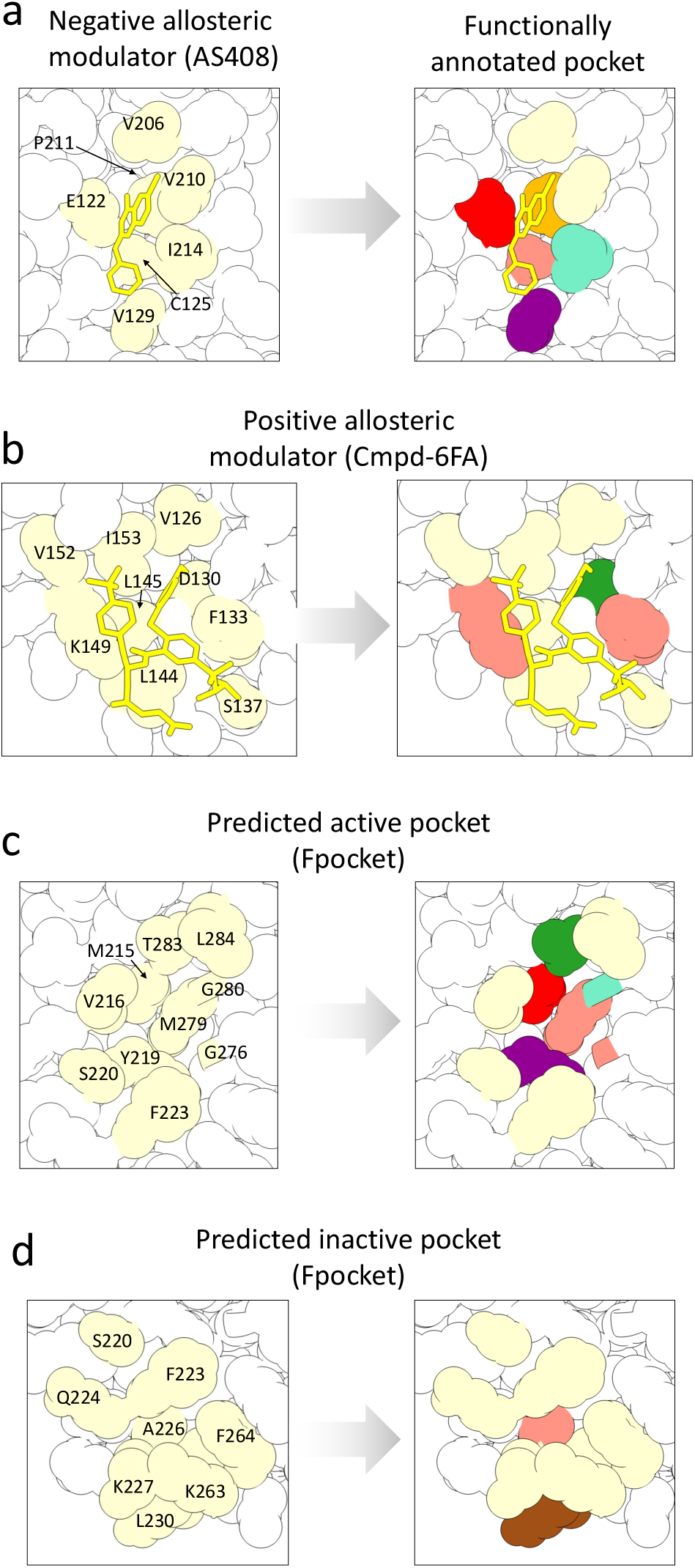
Rationalization and prioritization of allosteric pockets. **a** The binding pocket of known negative allosteric modulator, AS408, shown in yellow. PDB ID: 6OBA. Residues making direct contact with the NAM are colored beige. **b** Expression normalized maximum recruitment through Nb80, recruitment bias, and expression normalized potency of the mutations at sites making up the binding pocket of AS408. Shown below the heatmaps are the categorical annotations of significant residues: purple, green, and red indicate LoF, GoF, and CoF residues for maximum recruitment. Orange, blue, and magenta indicate Nb80 biased, β-Arrestin, biased, or bias switches. Aquamarine, brown, and salmon indicate LoF, GoF, and CoF for potency. **c** The binding pocket of AS408 with annotated residues colored. For residues annotated in both maximum recruitment and potency analyses, the color from maximum recruitment was used. **d, e, f** Same as in (a, b, c) but for the positive allosteric modulator Cmpd-6FA, with structure 6N48. **g, h, i** Same as in (a, b, c) but for a predicted active pocket in structure 2RH1. **j, k, l** Same as in (a, b, c) but for a predicted inactive pocket in structure 2RH1.

While the AS408 pocket is replete with functional residues, Cmpd-6, a positive allosteric modulator^72^ (PAM), binds in a pocket formed by 9 residues between TMs 2, 3, 4, and ICL2 (Figure 7b) with only 4 annotations. However, one of the residues is D130^3x49^, where mutations cause strong GoF for both maximum recruitment and EC_50_ (Supplementary Figure 7b), potentially explaining the PAM activity of Cmpd-6. Additionally, while Cpmd-6 mostly forms hydrophobic interactions with β2AR, it does form one hydrogen bond with K149^4x41^, which is an EC_50_ CoF residue. Taken together, our functional annotations are consistent with the activities of the two known β2AR allosteric modulators.

To explore prospective functional annotation, we used Fpocket^73^ to predict pockets in the inactive β2AR (PDB ID:2RH1). One additional pocket, between TMs 5 and 6 (Figure 7c), has 8 functional annotations among 10 residues, with most being maximum recruitment or EC_50_ LoF or CoF, but also one GoF residue (Supplementary Figure 7c), suggesting a highly active pocket. Indeed, an allosteric agonist was recently found to bind and activate another GPCR, GPR88, at this same site^74^. On the other hand, a nearby and partially overlapping pocket also between TMs 5 and 6 (Figure 7d) has only two annotations among 9 residues (Supplementary Figure 7d), suggesting that perturbation of this pocket is less likely to have functional effects on receptor activation.

## Discussion

We have presented here GPCR-MAPS, a general, modular, and massively scalable approach that uses pooled experiments to precisely quantify the effects of mutations on the activity and bias of GPCRs. GPCR-MAPS makes multiple important technical advancements over existing technologies for multiplexed receptor assessment^27–30^.

First, GPCR-MAPS directly quantifies the recruitment of proteins to receptors, so minimizing the confounding effects of non-linear and unequal signal amplification, and feedback by endogenous signaling networks. Second, GPCR-MAPS is highly modular - we demonstrate here the successful use of a nanobody, a mini-G protein, and β-arrestin as signal transducers. The receptor-proximal readout also facilitates comparisons across these signal transducers, which can otherwise be confounded by differential signal amplification and crosstalk^75^. This modular design allows direct study of GPCR bias at the level of recruitment to a receptor. To further demonstrate generalizability, we also show the vasopressin 2 receptor recruitment of miniGs and β-arrestin in response to the endogenous ligand arginine vasopressin (Supplementary Figure 6e). Third, GPCR-MAPS measures and quantitatively deconvolves the strong confounding influence of receptor expression changes on signaling parameters. Fourth, due to its highly multiplexed design and straightforward read out, GPCR-MAPS extends GPCR mutational scanning to the quantification of full dose-response curve activation functions for all possible mutations in a receptor. Comparing our data to previous β2AR mutagenesis studies reveals the importance of all four of these innovations (Supplementary Figure 1i-k, Figure 6b).

Applied to β2AR, GPCR-MAPS generates over >150,000 measurements and charts a complete amino acid resolution picture of how quantitative multi-modal receptor phenotypes are encoded and how they can be tuned. The β2AR maps demonstrate the importance of quantitatively controlling for receptor expression changes. After deconvolving mutational effects on expression and recruitment, the residues that disrupt recruitment form a contiguously connected core activation network linking the ligand binding site to the transducer coupling site (Figure 8a). The data also identify the extracellular vestibule, particularly ECL2, as a major allosteric site to control receptor bias, and TM7 near the NPxxY motif as an internal region that allosterically controls receptor activation bias (Figure 8b).

**Figure 8.**
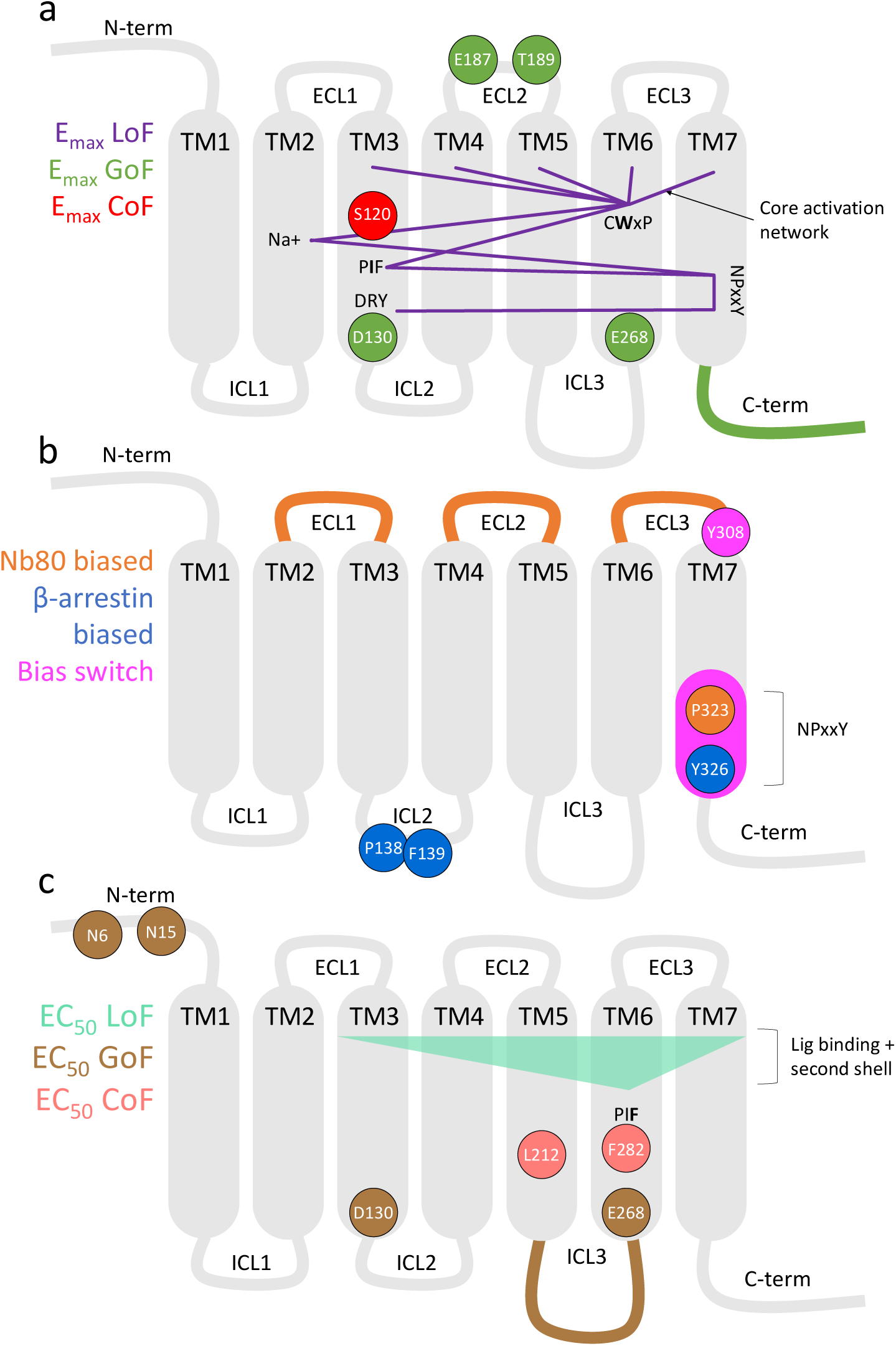
Summary of findings. **a** Selected findings from the maximum recruitment analysis. Core activation network shown as purple lines, residues or topological domains colored according to annotation. **b** Selected findings from biased recruitment analysis. NPxxY motif collectively illustrated as “bias switch” as there are multiple bias functional residues in and near this motif. **c** Selected findings from the potency analysis. Ligand binding and second shell residues illustrated as triangle.

Quantifying the full activation functions of >7,500 receptor variants in a single experiment (Supplementary Figure 8) revealed that expression is also a major influence on EC_50_. This has been previously predicted by theoretical^64–66^ models and agrees with recent work on plant hormone receptors^35^. After correcting for the influence of expression changes, EC_50_-modulating mutations are topologically localized, including potency increasing variants in ICL3 and at both N-glycosylation sites in the unstructured N-terminus (Figure 8c). Mutations that decrease potency are in close proximity to the core activation network and so likely modulate it. In contrast, residues increasing potency are far from the core activation network, likely affecting potency via mechanisms independent of the core activation network. This distinction in the location and likely mechanism of LoF and GoF variants parallels observations in the Src kinase, where mutations that decrease kinase activity are enriched close to the active site but mutations that increase it are not^76^.

GPCRs are the largest family of membrane receptors and the most successful class of drug targets, with one third of current clinical therapeutics targeting GPCRs^7^. Precision modalities such as biased signaling and allosteric modulation could lead to GPCR drugs with improved efficacy and safety, but these drugs have historically only been found by serendipity. Despite a revolution in GPCR structural biology, we lack sufficient functional understanding of GPCR activation and effector recruitment to design biased or allosteric drugs. We believe that quantifying multi-modal mutational effects will allow the high-resolution functional annotation of many human receptors and accelerate the development of new precision drugs.

Residues biased toward Nb80 recruitment are enriched in ECL2, which forms part of the exosite of salmeterol, a β2AR agonist strongly biased toward G protein signaling^59^, suggesting mutant behavior is related to drug activity binding at the same sites. Likewise, known allosteric modulators bind in pockets with high impact mutations that match their activities. AS408, a NAM, binds in a pocket with loss- and change-of-function residues, while Cmpd-6FA, a PAM, binds in a pocket with gain- and change-of-function residues. In addition to these known allosteric sites, GPCR-MAPS prioritizes an additional – and, to our knowledge, novel – allosteric pocket to target in β2AR and deprioritizes one where mutations only have limited effects.

The GPCR-MAPS approach is modular and scalable, and its application across the GPCR superfamily will enable a better understanding of the largest family of human receptors. To date GPCR drug discovery has proceeded from an era of blind high-throughput screening to an era of structure-based drug discovery. We believe we are now entering a third era of GPCR drug discovery, where high-resolution functional maps empower the rational design of precision GPCR therapeutics, including highly specific biased and allosteric modulators.

## Materials and Methods

### Plasmids

Plasmids encoding SPARK2 components were gifts from Christina Kim and Alice Ting and were obtained from Addgene (Addgene plasmid # 125228, 104839, 104840, 104841). From these plasmids we generated new plasmids using Gibson cloning. attB-HA-b2AR-eLOV-TEVcs-Gal4-UAS-Citrine encodes the receptor fusion followed by the bGH poly(A) signal, then the UAS sequence followed by Citrine fluorescent protein. A 26 nucleotide barcode was inserted in the 3’ UTR of the Citrine sequence. To make cloning more efficient, we transferred the fusion coding sequence from the NanoLuc-15aa linker-βarrestin2-HA-GS linker-TEVp plasmid from the pAAV vector (which contains GC-rich inverted terminal repeats) to the pUC19 vector. We then used site directed mutagenesis to introduce the S153N variant to TEV, generating uTEV1Δ). We then made versions of this plasmid with Nb80 and miniG_s_ sequence instead of β-Arrestin. We also modified the miniG_s_ vector so that uTEV1Δ was fused N-terminally (NanoLuc-15aa linker-GS linker-TEVp-miniG_s_). pCAG-NLS-HA-Bxb1 was a gift from Kenny Matreyek and Doug Fowler.

### Development of GPCR-MAPS recruitment assays

We started with the SPARK2 system^41^ and extensively re-engineered the system to enable massively multiplexed experiments. First, we combined the GPCR construct and the reporter construct onto a single plasmid and added a DNA barcode (26 nucleotides) in the 3’ untranslated region (UTR) of the Citrine reporter so that the GPCR variant on any given plasmid can be identified (Figure 1a and Supplementary Figure 1e). We also added the attB recombination site to enable integration into HEK293T landing pad cells^40^, which ensures copy number of exactly one in each cell. We used the active-state-specific nanobody Nb80^42^ as the binding partner to optimize the assay. While the original assay used transient transfection to introduce transgenic material and took place over the course of minutes, we found that substantially increased times were necessary to obtain detectable signal, presumably due to the reduction of copy number of the receptor and reporter constructs (24 hours, Supplementary Figure 1a). Further, we tested evolved variants of both TEV and the LOV domain, uTEV1Δ^77^ and hLOV1^78^; the combination that yielded the highest signal to noise ratio was uTEV1Δ and eLOV (Supplementary Figure 1b). Next, we also validated the use of β-arrestin and miniG_s_^79^ as the binding partner in this system (Supplementary Figure 1b). Finally, in order to account for differences in barcode abundance in the library, as well as barcode-specific effects on RNA abundance, we also include a condition in which Gal4 is expressed alone (Supplementary Figure 1f); this condition should maximally stimulate all barcodes and provides normalization for the experimental conditions^27^.

### Saturation mutagenesis library creation

We designed six pools of mutation-bearing oligonucleotides, each pool representing mutated coding sequence for 66-70 amino acids of the full-length (413 amino acids) b2AR with flanking wild-type sequence for amplification from the pool (using primers dialout_tile[1-6]_[F/R], Supplementary Table 5). For each pool we designed primers to amplify the whole plasmid except for the mutated segment (using primers b2AR_satmut_tile[1-6]_[F/R]), then used Gibson assembly to introduce the mutagenic oligos. Oligos were designed to encode all possible single amino acid changes (using the human optimized codon for each alternative amino acid), all premature stop codons, and all synonymous codons when possible (using the most optimal codon, excluding the wild-type codon).

### Barcode integration and variant-barcode association

10 μg of the attB-HA-b2AR-eLOV-TEVcs-Gal4-UAS-Citrine saturation mutagenesis plasmid was digested with 40 units PpuMI (New England Biolabs) overnight at 37°, then in the morning 10 units more PpuMI were added and 30 minutes more incubation at 37°. Then the cut band was gel purified. The barcode construct (oTLM_257) was made double stranded by PCR amplifying with oTLM_258 and oTLM_259 (Supplementary Table 5). Double stranded barcode was incorporated into the cut plasmid backbone by Gibson assembly and transformed into electrocompetent bacteria. Following recovery in SOC media, we plated 0.1% of the transformation to estimate the number of clones, and inoculated the rest into LB media with ampicillin for overnight growth. Estimated number of clones was 300,000. In the morning the bacteria was pelleted and plasmid was midi-prepped (Qiagen Plasmid Plus Midi Kit).

We used PacBio long read sequencing to associate each variant with the barcode. We digested the plasmid pool with BsiWI (New England Biolabs) to release the coding sequence and barcode and SMRT bell adapters were ligated according to manufacturer’s instructions. The library was sequenced on a PacBio Sequel IIe. We used alignparse^80^ to align reads, quality filter, and call sequences for each barcode (Supplementary Figure 1).

### Landing pad integration

HEK 293T LLP-iCasp9-Blast^40^ cells were a gift from Kenny Matreyek. Cells were grown in DMEM with GlutaMAX (Gibco), with Tetracyline-free fetal bovine serum. 14 million cells were plated onto a T225 flask. The next day, transfection was done with Lipofectamine 3000 (Invitrogen) with 17.5 μg of pCAG-NLS-HA-Bxb1 and 17.5 μg of the mutated and barcoded attB-HA-b2AR-eLOV-TEVcs-Gal4-UAS-Citrine. The cells were allowed to incubate for two days. Then, the media was removed and replaced with media containing 2 μg/mL doxycycline and incubated for one day. Then, the media was changed for media containing 2 μg/mL doxycycline and 10 nM rimiducid (MedChemExpress) and incubated for one day. Then media was changed to include just 2 μg/mL doxycycline and outgrowth of cells was allowed for ∼one week, passaging cells when they reached 90% confluency.

### GPCR-MAPS recruitment experiment

10 million cells were plated onto each T175 flask and allowed to incubate overnight. The next day, the cells were transfected with 36 μg of transducer plasmids (pUC19-NanoLuc-β-arrestin-uTEVp1 or pUC19-NanoLuc-Nb80-uTEVp1), using Lipofectamine 3000 and allowed to incubate overnight. The next morning, cells were treated with media containing furimazine and either drug or DMSO: [13.05 mL DMEM, 126.56 μL furimazine, 270 μL 1M HEPES, and 13.5 μL isoproterenol or DMSO]. Furimazine was obtained from the Nano-Glo Live Cell Assay System (Promega). Isoprenaline was obtained from Sigma (catalog number 15627). Cells were incubated in the dark (covered with aluminum foil) at 37° for 24 hours. Then, cells were lifted with Trypsin-EDTA, pelleted and stored in -80 until RNA extraction. For the Gal4 maximum stimulation condition, after 10 million cells were plated and grown overnight, 36 μg of pUC19-CMV-Gal4 was transfected with Lipofectamine 3000. The cells were allowed to incubate for 30 hours, then cells were pelleted and stored at -80°.

### RNA isolation and sequencing library preparation

For the first set of experiments (Nb80 and β-arrestin recruitment with 10 μM isoprenaline), cell pellets were thawed and resuspended in buffer RLT with added beta-mercaptoethanol. Then, they were homogenized by centrifuging through QIAshredder columns. RNA was then purified from the homogenate with RNeasy Midi-prep kit (Qiagen) and eluted with 400 μL of water. Between 120 and 160 μg of total RNA was used as input for reverse transcription (RT) reactions (6-8 μg per reaction, 20 reactions per sample) using SuperScript III reverse transcriptase (Invitrogen). For the dose response experiment, Dynabeads mRNA direct purification kit (ThermoFisher Scientific) was used according to manufacturers instructions and mRNA was eluted in 80 μL of Tris-HCl. Total mRNA yields were between 11.5 μg and 16 μg. Between 5.75 and 8 μg mRNA was used as input for RT (1.15-1.6 μg per reaction, 10 reactions per sample). For both sets of reactions, RT was carried out with 2 pmol of primer (oTLM_266, Supplementary Table 5) and incubated at 55° for one hour then 75° for 15 minutes. Following this, RNase A was added to a final concentration of 100 μg/mL and reactions incubated at 37° for 30 minutes. Then, reactions were cleaned up with Nucleospin PCR cleanup columns (Macherey Nagel) and eluted in 50 μL of elution buffer. 0.5 μL of the cDNA eluate was used in a qPCR reaction (Q5 High-Fidelity polymerase, New England Biolabs, and EvaGreen fluorescent dye, Biotium) to determine the number of cycles required to reach near-plateau phase. Then, 40 μL of cDNA was amplified across 4x 50 μL PCR reactions with the program 98° for 30s, followed by 13-16 cycles of [98° for 15s, 64° for 30s, 72° for 30s].

Reactions were column purified and 4% of this was used as input for a second round PCR, which added the rest of the Illumina adapter as well as unique index sequences. The second round PCR was 98° for 30s, followed by 5 cycles of [98° for 15s, 64° for 30s, 72° for 30s]. Frameshifting primers were used to introduce variability in the first few cycles of the sequencing reaction (oTLM_277-282, Supplementary Table 5). Reactions were then gel purified and analyzed on TapeStation (Agilent Technologies). Sequencing was done with Illumina NovaSeq with 2x50 paired end reads.

An additional step was taken for the dose response experiment. Following RT, we performed triplicate qPCR reactions with each cDNA sample, along with a set of serial dilutions of the original library plasmid, in order to estimate the relative quantity of Citrine RNA produced from each experiment. We used primers oTLM_021 and oTLM_024, which amplify the 5’ UTR of Citrine. We chose to amplify this region instead of the barcode (as was done for the library prep) to avoid idiosyncratic effects of amplification efficiency caused by diverse and potentially different barcode identities across different samples. Following qPCR we fit a standard curve (Supplementary Fig. 5b) then used this to estimate RNA abundance for the dose response samples (Figure 5b).

### Surface expression experiment cell culture and FACS

Cells were cultured in 245x245mm cell culture flasks until ∼80% confluent. Then, cells were lifted with Trypsin-EDTA, pelleted, supernatant removed, and resuspended in blocking buffer (1% bovine serum albumin in phosphate buffered saline). Cells were counted and 50 M cells were transferred to a 5 mL conical tube with 3.5 mL of blocking buffer, and incubated on ice, covered from light, with rocking for 30 minutes. Then, anti-HA antibody (Cell Signaling HA-Tag 6E2 Alexa Fluor 647 Conjugate) was added at dilution of 1:100 and incubated on ice, covered from light, with rocking for 60 more minutes. Then, cells were pelleted, supernatant removed, and resuspended in blocking buffer. Sorting was done on the BD FACSAria instrument. Bins were drawn so that each represented ∼25% of cells. In total, ∼20M cells were sorted for replicate 1, and ∼30M cells were sorted for replicates 2 and 3. 30-45M 2x50 basepair reads were collected on an Illumina NovaSeq instrument for each replicate.

### Surface expression sequencing library preparation

Sorted cells were pelleted and DNA extracted using Qiagen DNeasy Blood & Tissue kit, using 2 columns for each sample. 200 μL elution buffer was used to elute from each column, then eluates for each sample were combined. Then, 320 μL of each sample was was used as input for 16x 50 μL PCRs with primers adjacent to the barcode that include partial Illumina adapters and also varying lengths of degenerate bases to increase nucleotide complexity in the first few cycles of the sequencing run (oTLM277-oTLM282, Supplementary Table 5). Cycling was 98° for 30s, followed by 25 cycles of [98° for 15s, 64° for 30s, 72° for 30s]. Reactions were column purified and 4% of this was used as input for a second round PCR, which added the rest of the Illumina adapter as well as unique index sequences. The second round PCR was 98° for 30s, followed by 5 cycles of [98° for 15s, 64° for 30s, 72° for 30s]. The products of the second PCR were gel purified and analyzed on TapeStation, then sequenced on Illumina NovaSeq with 2x50 paired end reads.

### Sequencing data processing

Paired-end sequencing reads were first merged with vsearch^81^ and then adapters were trimmed with cutadapt^82^. This leaves each read representing the barcode sequence, which was then matched with barcodes identified in the variant-barcode association with a custom script.

### Calculation of surface expression scores

Surface expression scores were calculated as previously described^25^. Briefly, the frequency of each variant in each bin is enumerated by combining counts of all barcodes associated with that variant. The frequency, along with the total number of cells sorted into each bin, is used to estimate the number of cells in each bin. Then, the log10 transformed geometric mean fluorescence value of all cells sorted into each bin is combined with the estimated cell counts to arrive at the raw surface expression estimate. These estimates were then normalized between 0 (represented by the median of presumed loss-of-function variants: e.g. premature stop codons before the 300^th^ amino acid) and 1, the wild-type score. Surface expression scores were considered high confidence if the estimated number of cells sorted was >50.

### Calculation of recruitment scores

First, barcode read counts for all barcodes representing the same variant were combined and then the frequency of that variant was calculated by dividing that number by the total number of barcode counts in the sample. Then, to arrive at a raw score, we divided the variant frequency in the experimental sample by the variant frequency in the Gal4 sample, and take the log_2_ of this. The Gal4 sample represents a maximum stimulation condition in which the barcode frequency in the library, as well as any idiosyncratic effects of the barcode sequence on RNA stability should be captured. Raw scores are then scaled between 0 and 1, with the median of nonsense mutations set as 0 and the wild-type score set as 1. Recruitment scores were considered high confidence if the frequency of that variant in the Gal4 condition was >2x10^-5^.

### LOWESS modeling and identifying outlier positions

The python package statsmodels.nonparametric.smoothers_lowess^83^ was used with the following parameter values: frac=0.5, it=3, delta=0.0. Following curve fitting, residuals were calculated as the distance in the y-axis between the data point and the line, and z-scores were calculated as the residual / standard error of the mean. In the case of Nb80 versus β-arrestin, the average standard error of the mean between the two measurements was used.

### Structural analysis

Residue contacts were defined by GetContacts (https://getcontacts.github.io/index.html). We used the definition of the isoprenaline binding site as presented in the publication solving and describing the structure^44^. Specific interactions between residues and isoprenaline atoms were taken from the publication as well as GetContacts. The binding interface of β2AR with Nb80 was defined by GetContacts analysis of structure. The predicted inactive and active structures in Figure 6k were taken from GPCRdb and were produced with AlphaFold-Multistate^45^. These structures were used to enable visualization of the entire ICL3, which is not resolved in experimentally solved structures.

### Dose response curve fitting

In the package neutcurve^84^, we used the neutcurve.hillcurve.HillCurve function to fit curves with the following options: infectivity_or_neutralized=’neutralized’, fix_slope_first=True, fs_stderr=None, fixbottom=False, fixtop=False, fixslope=False, fitlogc=False, use_stderr_for_fit=False, no_curve_fit_first=False. Following the fitting procedure, we then filtered curves based on heuristics: R^2^ > 0.9, EC50 within the measured range (i.e. between 10pM and 10uM), and E_max_ < 1.5x the maximum measured recruitment value. See Supplementary Figure 5e for the number of variants filtered.

### Fpocket pocket prediction

We used PDB ID 2RH1 as the structural template for pocket prediction and used the Fpocket web server with default settings to predict pockets. We chose the two pockets to illustrate simply based on having many or few annotated residues.

## Acknowledgments

We thank Christina Kim for advice regarding the SPARK2 system. We thank the CRG/UPF Flow Cytometry Unit for technical assistance. We thank Jana Selent and lab members for helpful discussion. We thank all members of the Lehner lab for helpful discussion.

## Data availability

Files needed to reproduce analyses can be found at zenodo (https://zenodo.org/records/15363336). Raw sequencing reads can be found at Sequence Read Archive (accession number PRJNA1260170).

## Code availability

Custom code to reproduce analyses can be found at github (https://github.com/lehner-lab/GPCR-MAPS).

