## Supplementary figures for "GPCR-MAPS: high-resolution functional and allosteric mapping of G protein-coupled receptor activation and bias"

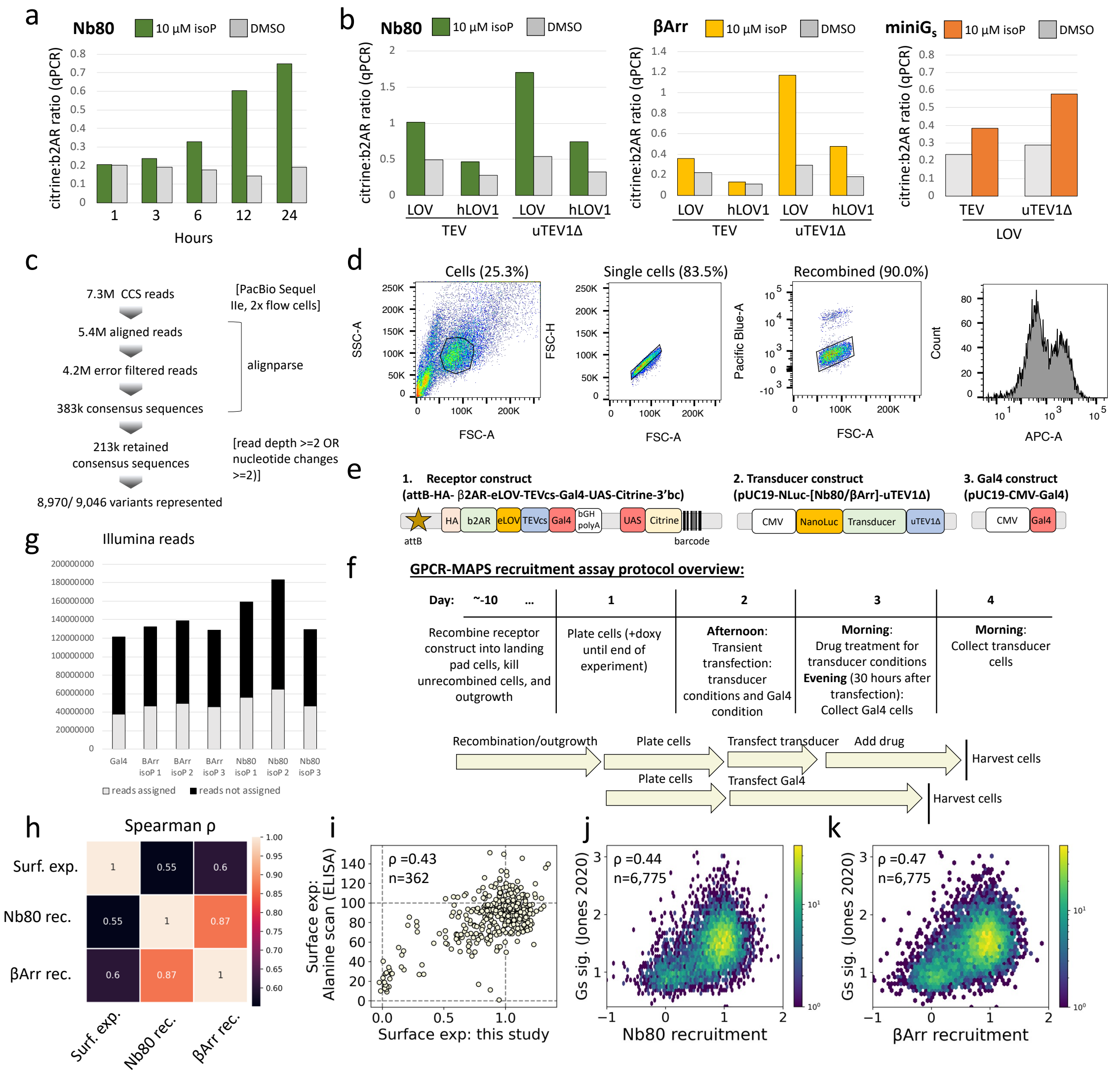

**Supplementary Fig. 1. GPCR-MAPS development**

**a** GPCR-MAPS timecourse experiment. Cells expressing the β2AR receptor construct in the landing pad were transiently transfected with transducer construct bearing Nb80 and the next day treated with 10 µM isoprenaline. Drug-treated cells were then incubated for varying amounts of time before cells were harvested and qPCR was used to measure reporter abundance. The qPCR was performed with a set of primers amplifying the Citrine reporter construct and a set of primers amplifying the β2AR coding sequence as a reference. Recruitment was then inferred by the ratio of Citrine to β2AR. **b** GPCR-MAPS experiment, performed with 24 hour drug incubation, but with different genotypes for the TEV protease and the LOV domain. Expriments were done as in (a) but with Nb80, β-Arrestin, or miniG$_s$ as the transducer. **c** Processing steps and numbers of reads/sequences at each step for the processing and analysis of PacBio long read sequencing linking variants to barcodes. **d** Gating strategy for FACS. **e** Schematics of the constructs used in GPCR-MAPS. The receptor construct encodes the HA-tagged receptor, fused to eLOV, TEVcs, and Gal4; downstream is Citrine driven by the UAS promoter with the barcode in the 3' UTR. This construct is integrated into the landing pad. The transducer construct contains a fusion of NanoLuciferase with the transducer and TEV protease, driven by CMV promoter. This construct is transiently transfected. The final construct is Gal4 alone, driven by CMV promoter. This construct is used to stimulate maximal reporter levels and serves to normalize differences in barcode abundance across the library as well as any barcode-specific effects on RNA stability in cells. **f** Timeline schematic of the GPCR-MAPS recruitment experiment, and the operations that are performed each day. **g** The number of Illumina reads acquired for each sample, with black and gray portions of bars indicating the fraction of reads that were assigned or not to barcodes. **h** Spearman correlations between surface expression, Nb80 recruitment, and β-Arrestin recruitment. **i** Surface expression measurements from this study compared to those generated in Heydenreich *et al*., 2023. **j** Nb80 recruitment versus $G_s$ signaling from Jones *et al*., 2020. **k** β-arrestin recruitment versus $G_s$ signaling from Jones *et al*., 2020.

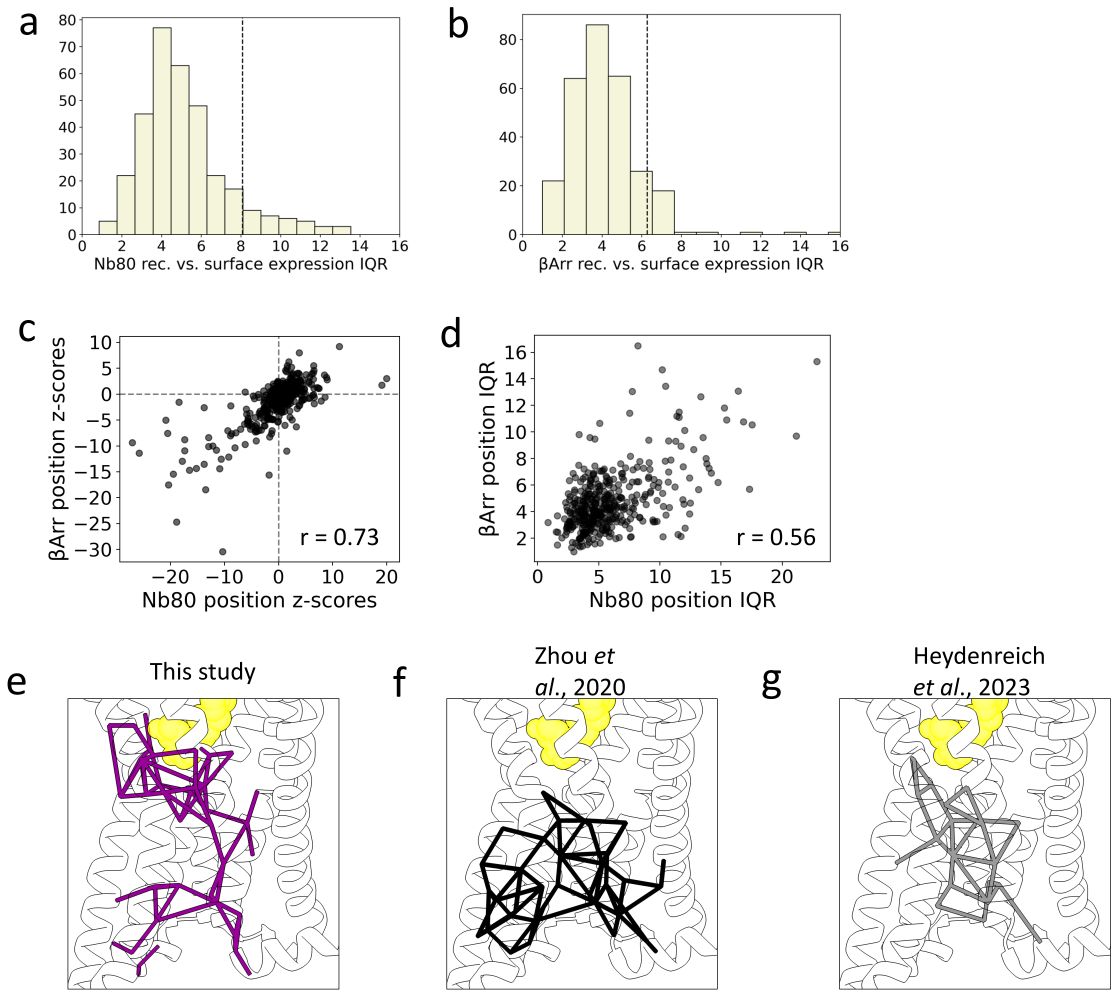


**Supplementary Fig. 2. Extended analysis of effects on recruitment beyond those on expression**

**a** Distribution of interquartile ranges of z-scores for all residues not in the loss-of-function or gain-of-function categories for Nb80 maximum recruitment. Dashed line indicates the $90^{th}$ percentile; residues to the right of this line are prioritized as change-of-function. **b** Same as in (a) but for β-Arrestin recruitment. **c** Position-average z-scores for Nb80 and β-Arrestin recruitment. **d** Position interquartile ranges of z-scores for Nb80 and β-Arrestin recruitment. **e** Activation network defined in this study, inter-residue contacts shown as purple bars. **f** Class A GPCR activation network defined by Zhou *et al*., 2020, inter-residue contacts shown as black bars. **g** β2AR activation network defined by Heydenreich *et al*., 2023, inter-residue contacts shown as gray bars.

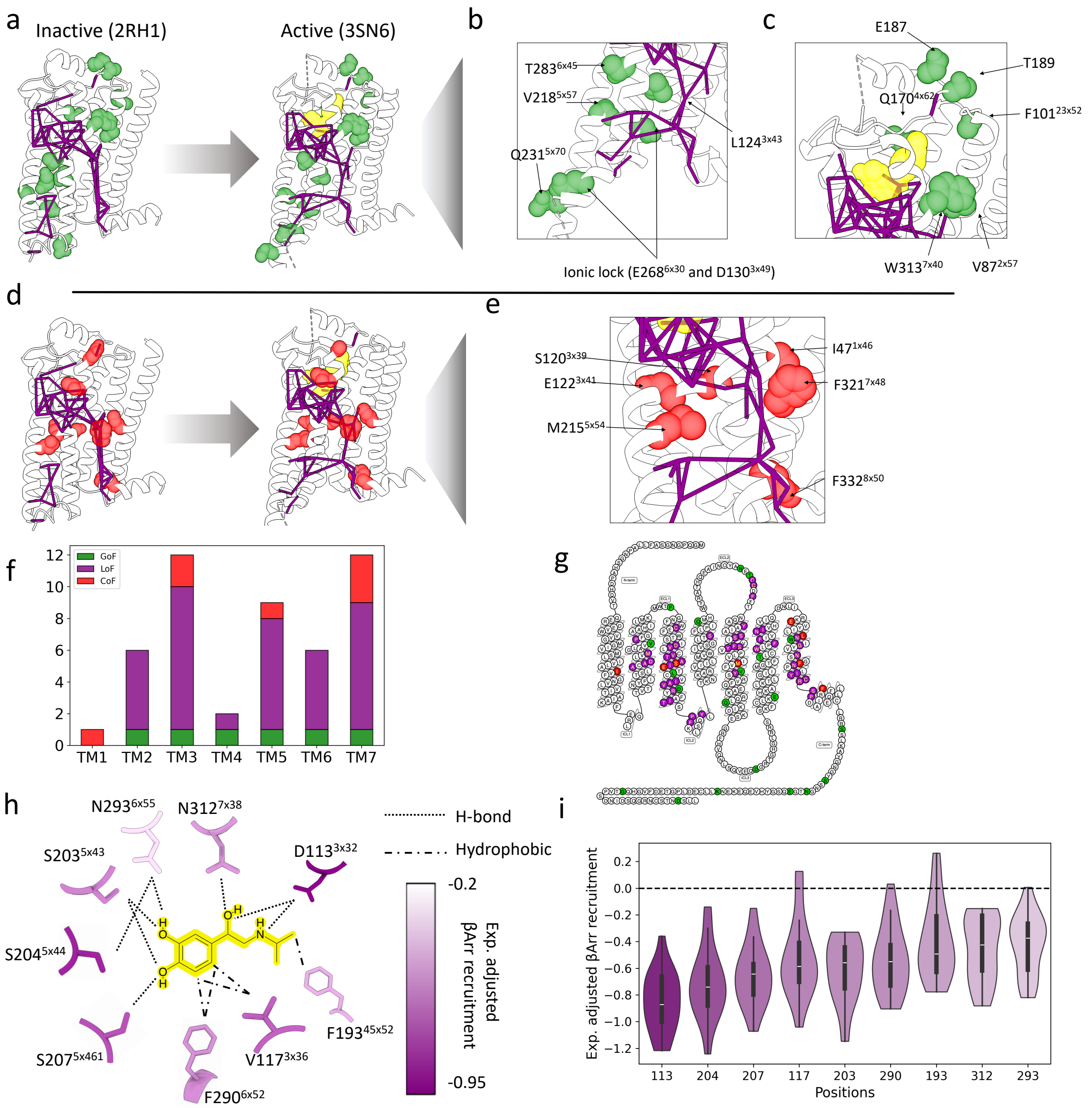


**Supplementary Fig. 3. Structural and sequence locations of maximum recruitment gain- and change-of-function residues.**

**a** Gain-of-function residues colored green on the inactive and active structures, with the core activation network represented as purple sticks. **b** Enlarged view of the ionic lock and surrounding residues in the active structure. **c** Enlarged view of the gain-of-function residues near the orthosteric site. **d** Change-of-function residues colored red on the inactive and active structures, with the core activation network represented as purple sticks. **e** Enlarged view of the change-of-function residues near the PIF and NPxxY motifs. **f** The number of each type of annotated residue in each transmembrane helix. **g** The sequence-based locations of the annotated residues in a snake plot representation (made with GPCRdb). **h** The isoprenaline binding pocket, with isoprenaline highlighted in yellow, and the residues making up the binding site colored according to their median expression adjusted β-arrestin recruitment level. **i** Violin plot of expression adjusted β-arrestin recruitment at the isoprenaline binding site residues. Violins are colored according to their median value, with the same color scale as in (h).

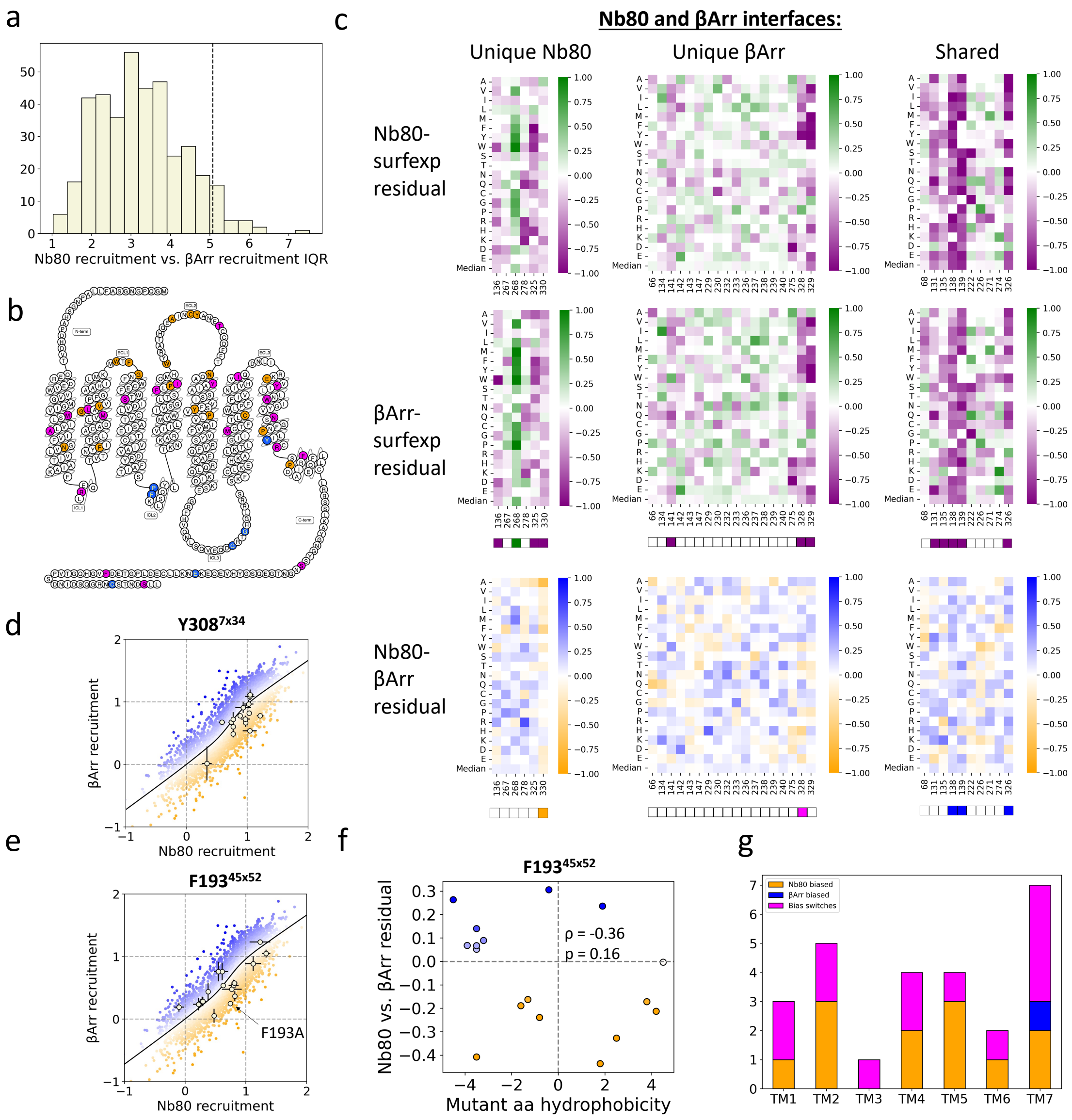


**Supplementary Fig. 4. Extended analysis of biased recruitment**

**a** Interquartile range of Nb80 to β-arrestin z-scores for all residues not called as Nb80 or β-Arrestin-biased. Dashed line indicates the 95$^{th}$ percentile of IQRs; residues with IQR greater than this line are considered as change-of-function. **b** Snakeplot representation showing the location of Nb80-biased, β-Arrestin-biased, and bias switch residues. **c** Mutation effects in the binding interfaces between β2AR and Nb80 and β-Arrestin. Shown are expression adjusted recruitment effects through both transducers, split into unique Nb80, unique β-Arrestin, or residues shared in both binding interfaces. At the bottom are the biased recruitment residuals for those same groups. Beneath heatmaps are indicated the residues annotated as maximum recruitment LoF, GoF, or CoF (purple, green, or red), and residues annotated as Nb80 biased, β-Arrestin-biased, or bias switches (orange, blue, or magenta. **d** Mutation effects at Y308$^{7x34}$ shown, with the population shown in the background and colored according to residual from the LOWESS fit line. Error bars represent standard error of the mean. **e** Same as in (d) but for F193$^{45x52}$. **f** Hydrophobicity of the mutant amino acid compared to the Nb80 vs. β-Arrestin residual for mutations at position F193$^{45x52}$. **g** The number of each type of annotated residue in each transmembrane helix.

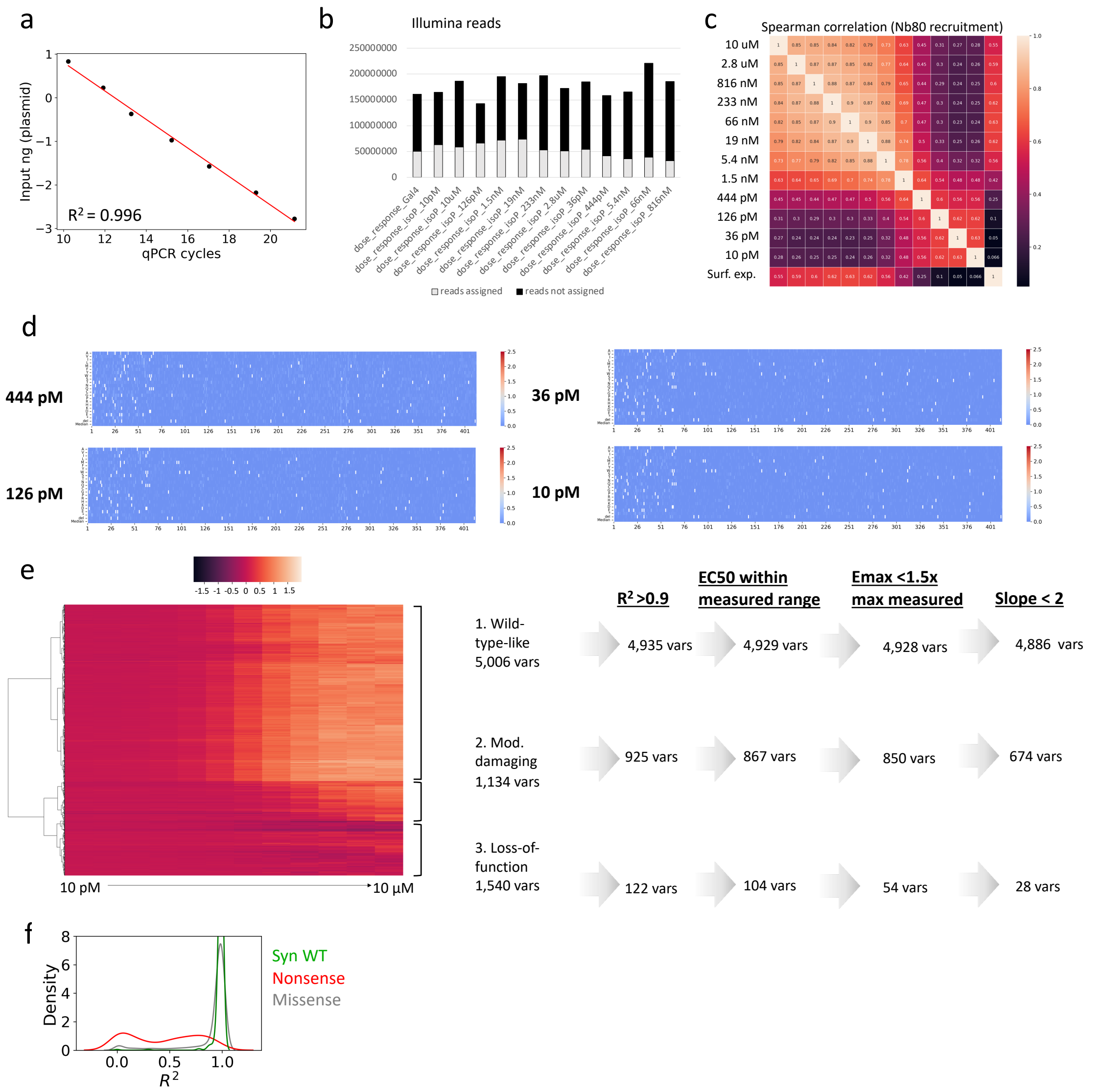


**Supplementary Fig. 5. Extended dose-response analysis**

**a** To measure RNA abundance, a standard curve was generated using serial dilutions of plasmid DNA and amplifying the Citrine reporter construct. **b** The number of Illumina reads acquired for each sample, with black and gray portions of bars indicating the fraction of reads that were assigned or not to barcodes. **c** Spearman correlations of variant scores in the serial dilution conditions or the surface expression experiment. **d** Heatmap representations of the recruitment data from the four lowest concentration conditions. **e** Clustered heatmap representations of the 7,680 measured dose response relationships. To the right are the quality control heuristics and the number of variants that pass each filter step from each cluster. **f** $R^2$ goodness-of-fit estimates for the curves fit to synonymous wild-type, nonsense, or missense variants.

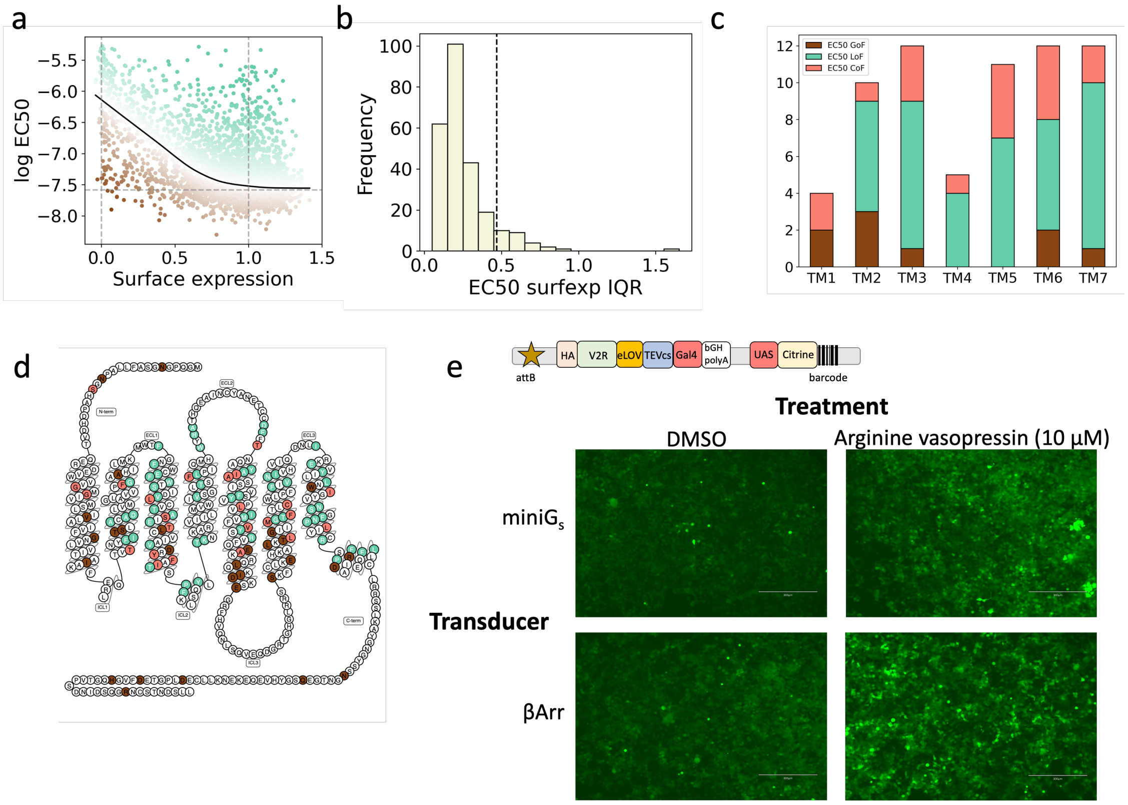


**Supplementary Fig. 6. Extended potency analysis**

**a** Log-transformed $EC_{50}$ compared with surface expression estimates, fit with a LOWESS regression and with variants colored by the residual to that line. **b** Distribution of position interquartile ranges of $EC_{50}$ to surface expression residuals for positions not identified as loss- or gain-of-function and with at least 10 variants with high quality dose response curves. **c** The number of each type of annotated residue in each transmembrane helix. **d** Snakeplot representation showing the location of $EC_{50}$ LoF, $EC_{50}$ GoF, and $EC_{50}$ CoF residues. **e** A library of vasopressin 2 receptor (V2R) variants were cloned into the same receptor construct as β2AR, then integrated into the landing pad, then cells were transiently transfected with either the miniGs or β-arrestin transducer constructs. Then, cells were treated with either DMSO or 10 μM arginine vasopressin, and 24 hours later the cells were imaged for expression of the fluorescent Citrine reporter. Scale bar is 300 μm.

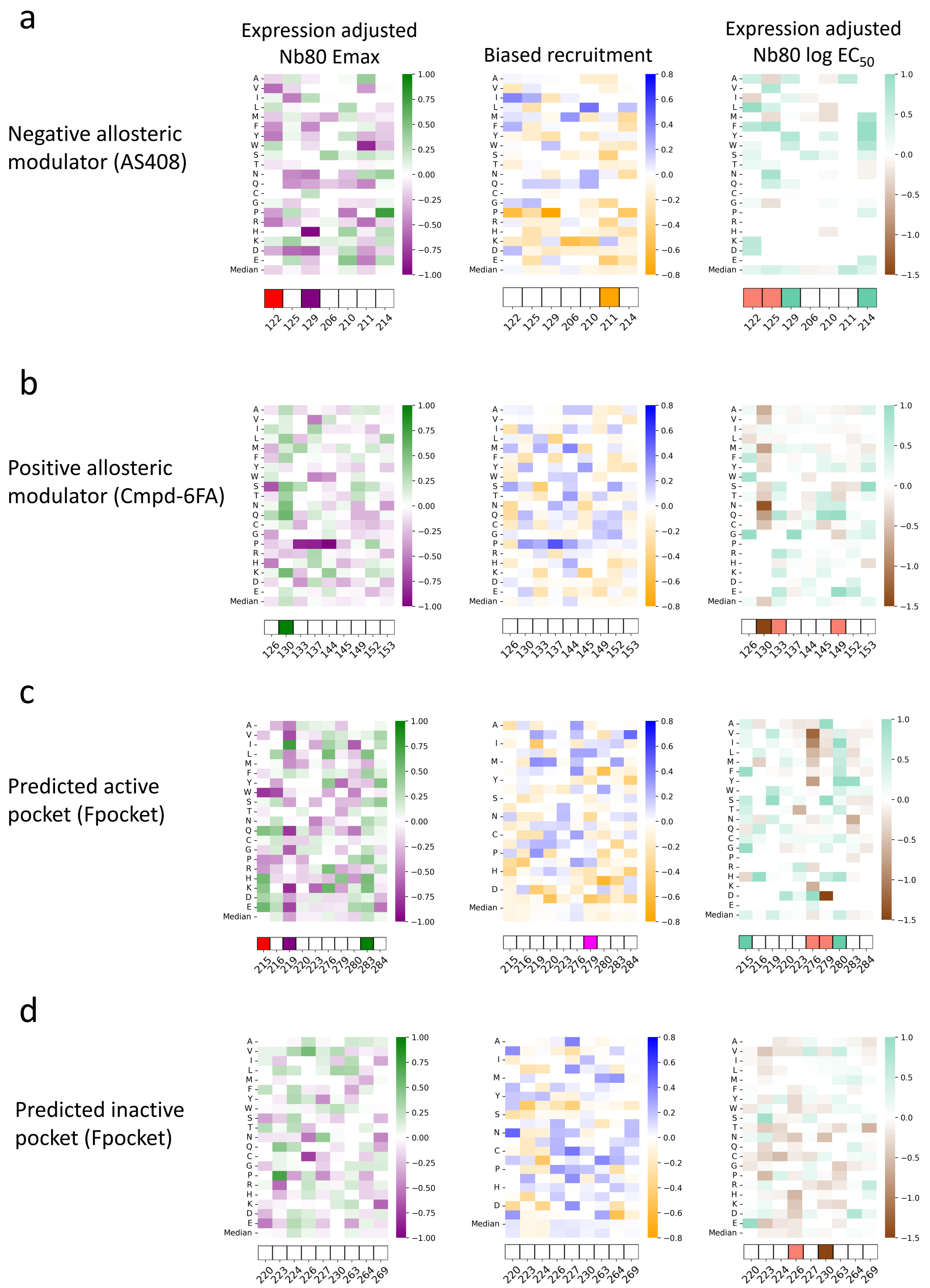


**Supplementary Fig. 7. Extended pocket annotation data**

**a** Expression adjusted maximum recruitment through Nb80, recruitment bias, and expression adjusted potency of the mutations at sites making up the binding pocket of AS408. Shown below the heatmaps are the categorical annotations of significant residues: purple, green, and red indicate LoF, GoF, and CoF residues for maximum recruitment. Orange, blue, and magenta indicate Nb80 biased, β-Arrestin, biased, or bias switches. Aquamarine, brown, and salmon indicate LoF, GoF, and CoF for potency. **b** Same as in (a) but for the positive allosteric modulator Cmpd-6FA, with structure 6N48. **c** Same as in (a) but for a predicted active pocket in structure 2RH1. **d** Same as in (a) but for a predicted inactive pocket in structure 2RH1.

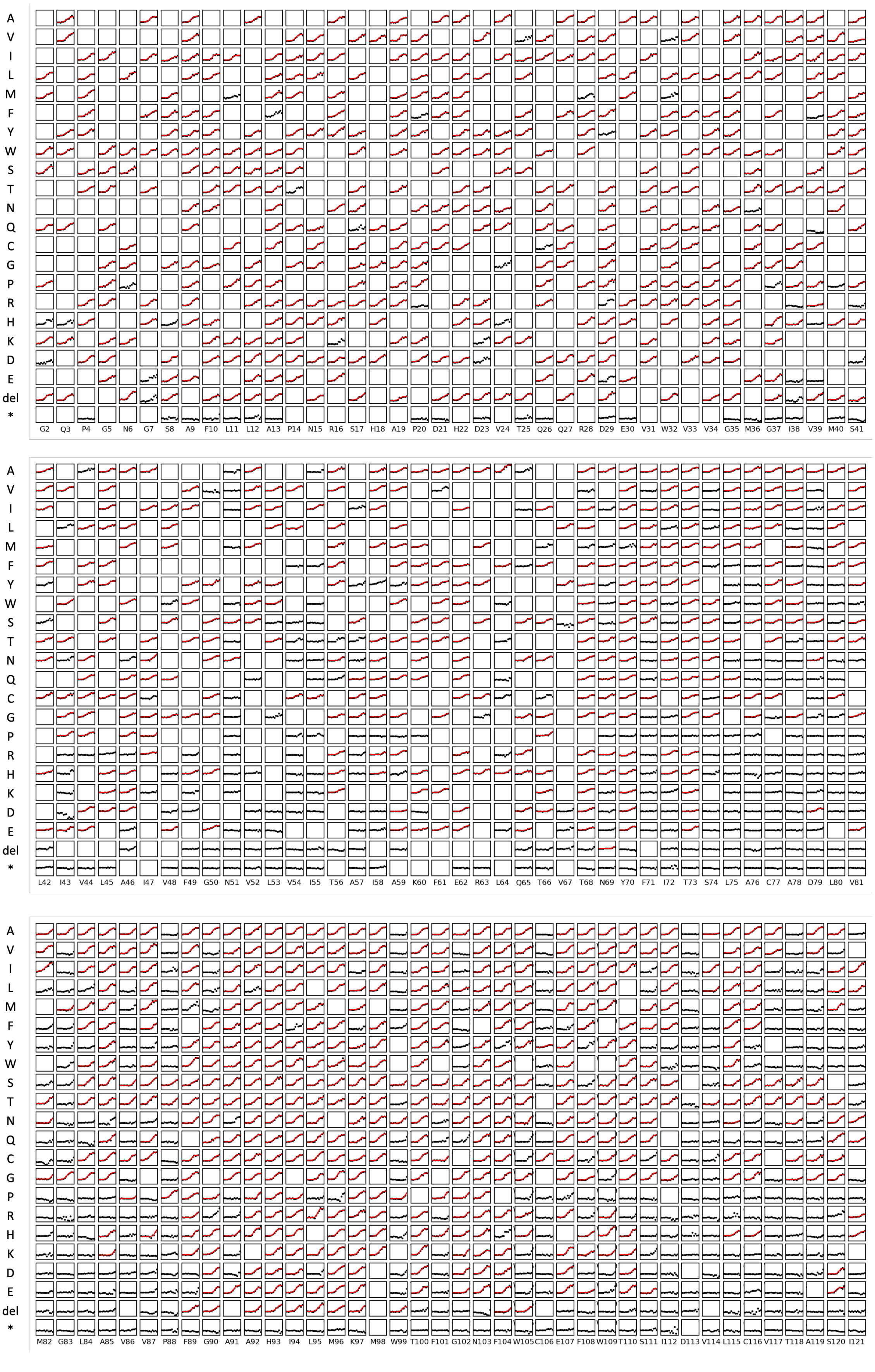

A V I L M F Y W S T N Q C G P R H K D E del *
G2 Q3 P4 G5 N6 G7 S8 A9 F10 L11 L12 A13 P14 N15 R16 S17 H18 A19 P20 D21 H22 D23 V24 T25 Q26 Q27 R28 D29 E30 V31 W32 V33 V34 G35 M36 G37 I38 V39 M40 S41
A V I L M F Y W S T N Q C G P R H K D E del *
L42 I43 V44 L45 A46 I47 V48 F49 G50 N51 V52 L53 V54 I55 T56 A57 I58 A59 K60 F61 E62 R63 L64 Q65 T66 V67 T68 N69 Y70 F71 I72 T73 S74 L75 A76 C77 A78 D79 L80 V81
A V I L M F Y W S T N Q C G P R H K D E del *
M82 G83 L84 A85 V86 V87 P88 F89 G90 A91 A92 H93 I94 L95 M96 K97 M98 W99 T100 F101 G102 N103 F104 W105 C106 E107 F108 W109 T110 S111 I112 D113 V114 L115 C116 V117 T118 A119 S120 I121

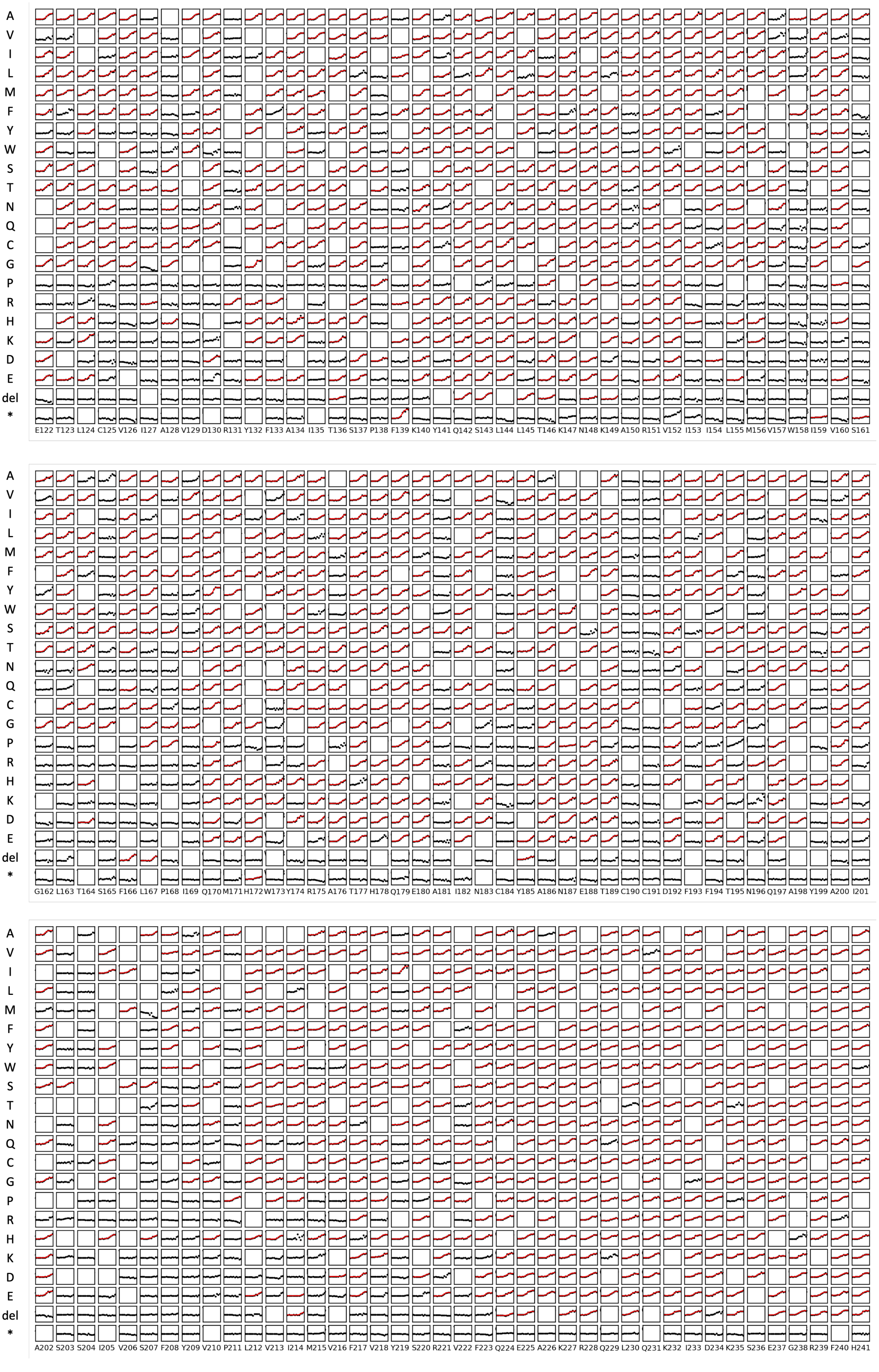
A V I L M F Y W S T N Q C G P R H K D E del *
E122 T123 L124 C125 V126 I127 A128 V129 D130 R131 Y132 F133 A134 I135 T136 S137 P138 F139 K140 Y141 Q142 S143 L144 L145 T146 K147 N148 K149 A150 R151 V152 I153 I154 L155 M156 V157 W158 I159 V160 S161
A V I L M F Y W S T N Q C G P R H K D E del *
G162 L163 T164 S165 F166 L167 P168 I169 Q170 M171 H172 W173 Y174 R175 A176 T177 H178 Q179 E180 A181 I182 N183 C184 Y185 A186 N187 E188 T189 C190 C191 D192 F193 F194 T195 N196 Q197 A198 Y199 A200 I201
A V I L M F Y W S T N Q C G P R H K D E del *
A202 S203 S204 I205 V206 S207 F208 Y209 V210 P211 L212 V213 I214 M215 V216 F217 V218 Y219 S220 R221 V222 F223 Q224 E225 A226 K227 R228 Q229 L230 Q231 K232 I233 D234 K235 S236 E237 G238 R239 F240 H241

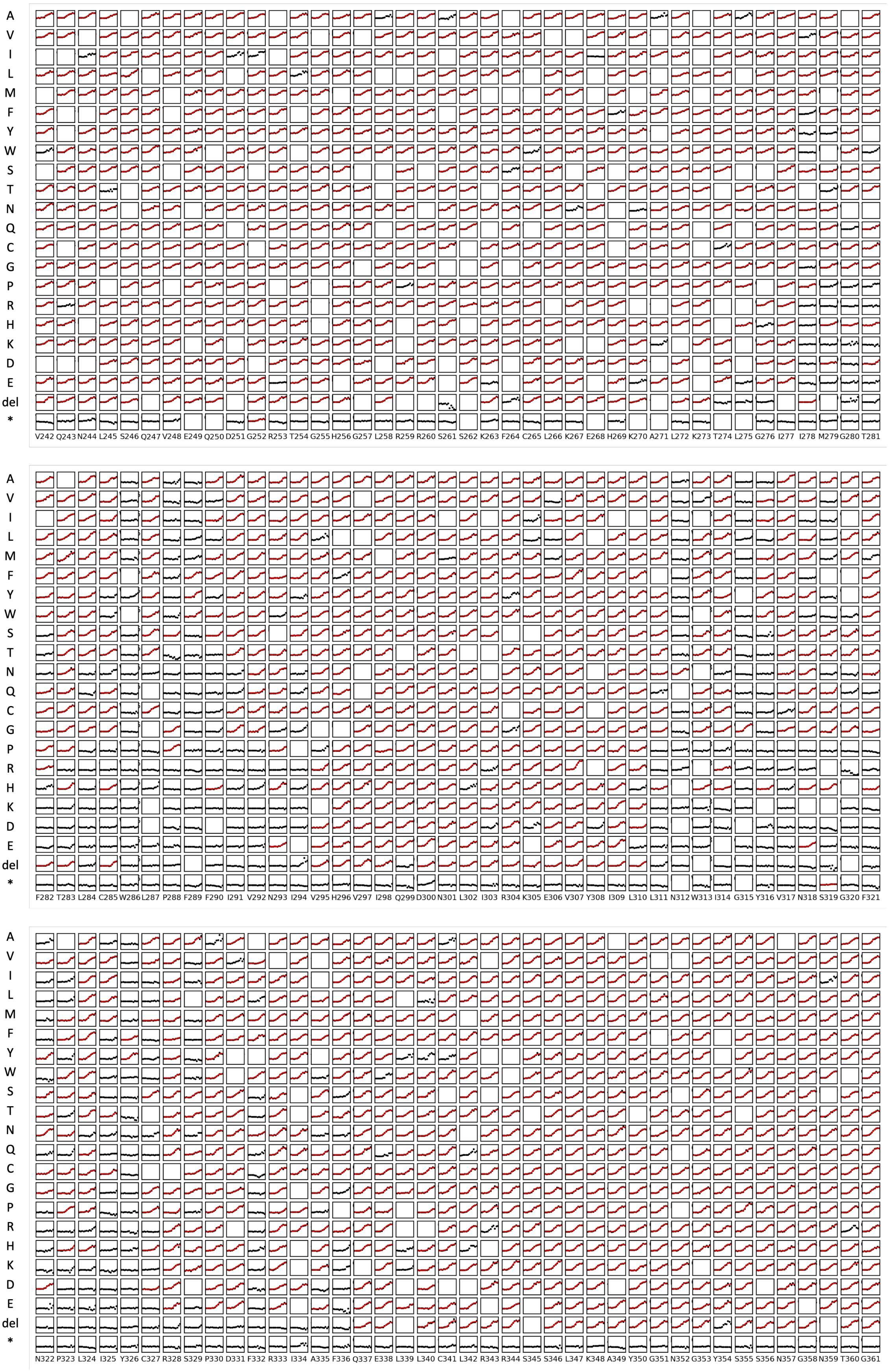
A V I L M F Y W S T N Q C G P R H K D E del *
V242 Q243 N244 L245 S246 Q247 V248 E249 Q250 D251 G252 R253 T254 G255 H256 G257 L258 R259 R260 S261 S262 K263 F264 C265 L266 K267 E268 H269 K270 A271 L272 K273 T274 L275 G276 I277 I278 M279 G280 T281
A V I L M F Y W S T N Q C G P R H K D E del *
F282 T283 L284 C285 W286 L287 P288 F289 F290 I291 V292 N293 I294 V295 H296 V297 I298 Q299 D300 N301 L302 I303 R304 K305 E306 V307 Y308 I309 L310 L311 N312 W313 I314 G315 Y316 V317 N318 S319 G320 F321
A V I L M F Y W S T N Q C G P R H K D E del *
N322 P323 L324 I325 Y326 C327 R328 S329 P330 D331 F332 R333 I334 A335 F336 Q337 E338 L339 L340 C341 L342 R343 R344 S345 S346 L347 K348 A349 Y350 G351 N352 G353 Y354 S355 S356 N357 G358 N359 T360 G361

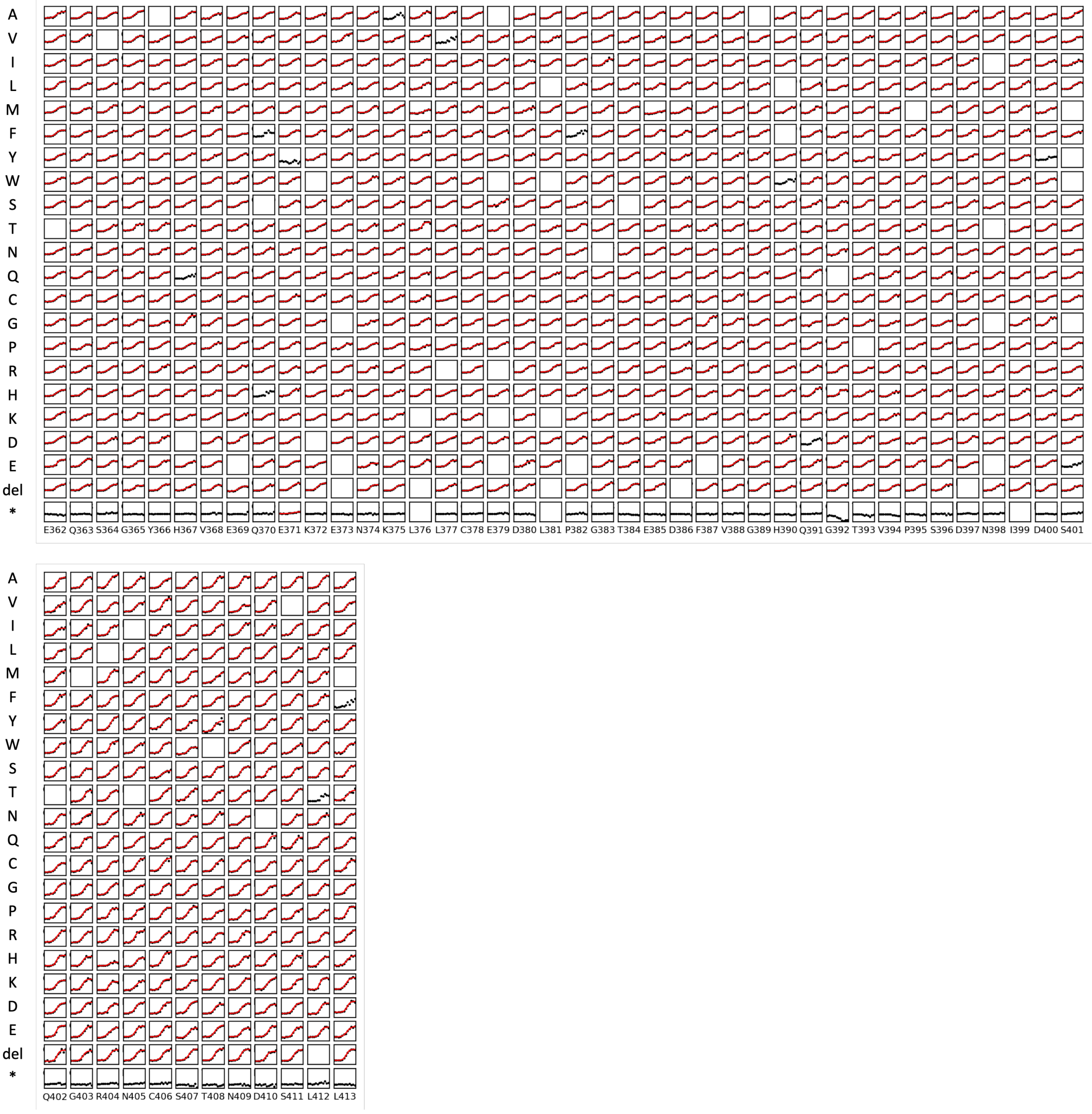


**Supplementary Fig. 8. Full set of dose response curves**

β2AR positions on the x-axis and variant identity on the y-axis. Cells without any data were filtered based on too low read count; cells with only black points passed the read-count filter but did not pass the curve fitting heuristic quality controls, cells with black points and red curves are the high-confidence curves.
